# Polarization and motility of one-dimensional multi-cellular trains

**DOI:** 10.1101/2023.07.02.547405

**Authors:** Jonathan E. Ron, Joseph d’Alesandro, Victor Cellerin, Raphael Voituriéz, Benoit Ladoux, Nir S. Gov

## Abstract

Collective cell migration, whereby cells adhere to form multi-cellular clusters that move as a single entity, play an important role in numerous biological processes, such as during development and cancer progression. Recent experimental work focused on migration of one-dimensional cellular clusters, confined to move along adhesive lanes, as a simple geometry in which to systematically study this complex system. One-dimensional migration also arises in the body when cells migrate along blood vessels, axonal projections and narrow cavities between tissues. We explore here the modes of one-dimensional migration of cellular clusters (“trains”), by implementing cell-cell interactions in a model of cell migration that contains a mechanism for spontaneous cell polarization. We go beyond simple phenomenological models of the cells as self-propelled particles, by having the internal polarization of each cell depend on its interactions with the neighboring cells, that directly affect the actin polymerization activity at the cell’s leading edges. Both Contact Inhibition of Locomotion (CIL) and Cryptic Lamellipodia (CL) interactions between neighboring cells are introduced. We find that this model predicts multiple motility modes of the cell trains, that can have several different speeds for the same polarization pattern. Comparing to experimental data we find that MDCK cells are poised along the transition region where CIL and CL roughly balance each other, where collective migration speed is most sensitive to the values of the cell-cell interaction strength.

## INTRODUCTION

Collective cell migration is essential for a variety of biological processes, from development to cancer progression [1]. The collective motion, in which several individual cells move as a single cohesive unit depends on the formation of the cell-cell contacts [2]. Upon the formation of cell-cell contacts, neighbouring cells affect each other at the contacts through chemical bonds that transmit forces [3, 4] and modify the polarization of the individual cells. One modification, termed Contact Inhibition of Locomotion (CIL) [5], denotes that colliding cells avoid moving in the direction of their neighbors, and stop or re-polarize away from the cell-cell contacts. Another modification of cellular migration due to cell-cell binding involves neighboring cells aligning their polarization, which eventually leads to a highly coordinated multi-cellular flow. Such polarization alignment occurs by forming Cryptic Lamellipodia (CL), actin-rich protrusions which are connected to the substrate and extend beneath the membrane of neighbouring cells [3, 6, 7].

While the role of cell-cell contact dynamics on cell migration has been extensively investigated in two dimensional experiments [5, 8–10], less is known about the role of cell-cell contacts on collective cell migration in one-dimensional environments. Confining cell migration to one-dimension greatly simplifies the system, by reducing its degrees of freedom, and enables to obtain more detailed insights into the mechanisms of cellular interactions, compared to two-dimensions. While most of the one-dimensional studies on cell-cell contacts focus on the CIL interaction between two colliding cells [11, 12], recent experimental work on one-dimensional cellular assemblies [13, 14] provides data regarding cellular interactions and coordination in cell trains, as well as data on the internal polarization of the cells [15]. In addition, one-dimensional systems allow for the development and analysis of tractable theoretical models. Theoretical analysis of these multi-cellular trains was mostly done so far in terms of self-propelled particles (SPP) models, where each cell can have a defined polarization direction that is affected by its neighbors [13, 16]. However, these models do not contain a mechanism for how cells polarize, and neglect their internal degrees of freedom.

Here we propose to go beyond previous models, and describe the migration of multi-cellular trains using a model which describes the mechanism of spontaneous internal polarization of each cell. For this purpose we utilize a simple model for one-dimensional cell migration of a cell with a fixed length, which has been verified by experiments and predicted the coupling between cell speed and persistence (UCSP), mediated by actin flows [17–19]. Within this model we directly incorporate the CIL and CL effects of the cell-cell contacts on the internal actin flow and actin-polymerization activity within the cells. We study the CIL and CL contact interactions in cell trains and the resulting collective migration patterns of cell trains, from cell pairs and up to long cell trains.

Using this model we explore the various global cell train migration regimes for different levels of noise in the actin-polymerization activity within the cells. For low noise levels we find co-existence of different stable cellular polarizations of the cells within the cell-train, which are beyond the current available models for collective cell migration that treat cells as SPPs [20–22]. We also show that when the noise level is relatively high two configurations are dominant, a persistent cell train where all cells are polarized in the same direction, and a stationary or slow moving train, where the edge cells are polarized away from each other. We compare the results to experiments conducted on MDCK epithelial cell pairs and triplets, in which the internal polarization of the cell is measured by a fluorescent marker of local actin polymerization activity.

## MODEL

In the experiments of MDCK cells migrating along fibronectin tracks (Fig.1A), the fluorescence intensity of the p21 Binding Domain of PAK fused to Yellow Fluorescence Protein (PBD-YFP) (Fig.1B) gives a measure of the local actin polymerization activity. This is a biosensor of active Rac1 and Cdc42, which both regulate actin polymerization near the membrane [16]. This measurement is used to determine the polarization of the cell, since where the signal is low/high we expect less/more actin polymerization activity, and thereby identify the back/front of the cell (Fig.1B). It is this internal cell polarization, as it is manifested in the competition between the actin polymerization activity at the two edges of the cell, that our theoretical model describes.

**FIG. 1:**
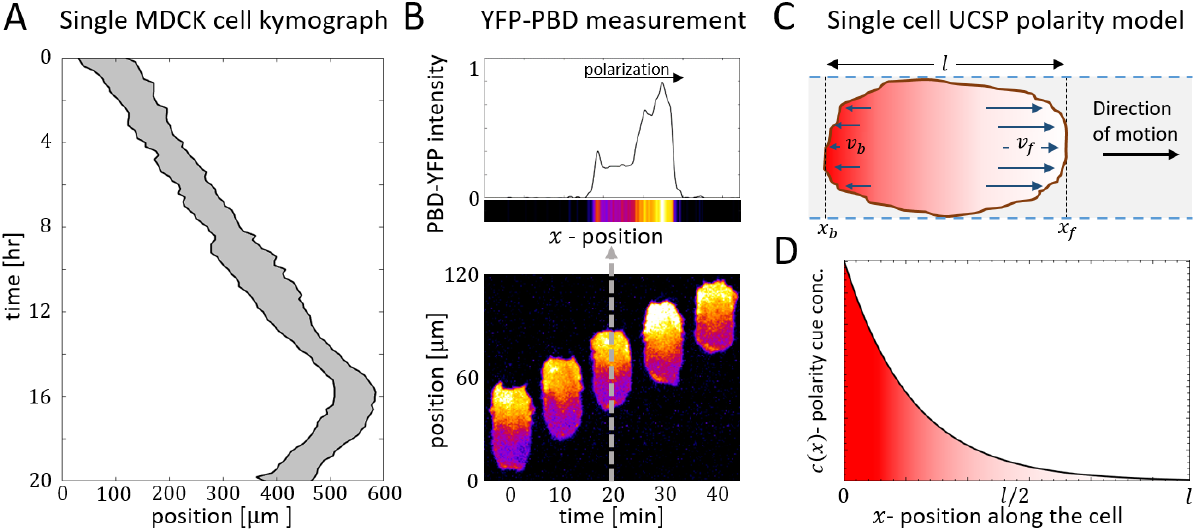
A) Kymograph of a single MDCK cell migrating along a fibronectin track. B) The YFP-PBD intensity measurement along a polarization axis of a single cell (Upper panel). Measurement corresponds to the dashed gray vertical line in the PBD-YFP cell kymograph in the lower panel. C) Illustration of the self-polarization cell model (UCSP): *x*_*f*_ and *x*_*b*_ denote the two edges of the cell. *v*_*f*_ and *v*_*b*_ denote the local polymerization speeds at *x*_*f*_ and *x*_*b*_ respectively. *l* is the length of the cell. Red/White color gradient indicates regions of high/low concentration of the polarity cue which inhibits actin polymerization at the edges of the cell. D) The concentration profile of the inhibitory polarity cue *c*(*x*) along the cell according to the model (Eq.2).

We present a model that describes the internal polarization of a one-dimensional cell, based on a model for cell polarization (UCSP model)[17]. Different extensions of the model were used to describe inhibitors/activators of actin polymerization in various types of cells in one-dimensional environments [18, 19, 23, 24]. To illustrate the model, consider a cell with a fixed length *l*, with the edges of the cell at *x*_*f*_ and *x*_*b*_ (front an back), and at each edge actin polymerizes at a mean speed of *v*_*f*_ and *v*_*b*_ respectively (Fig.1C). The net asymmetry in the retrograde flow across the cell length, can be simply quantified by the difference between local polymerization speeds

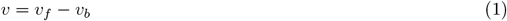

Consider now a polarity cue denoted by *c*(*x*), which inhibits actin polymerization at the edges of the cell near the membrane. Such inhibitory cues can be associated with capping proteins, which cap the edges of the polymerizing actin filaments and prevent their elongation [25], or proteins such as arpin which inhibit the formation of arp2/3 nucleation sites near the membrane [17, 26]. We assume that the inhibitory polarity cue is affected by the net asymmetry of the retrograde flow of the actin from the two edges of the cell (Eq.1) inside the cell. We assume that the coupling between the spatial distribution of the polarity cue to this net asymmetry of the retrograde flow rises due to an advection-diffusion process. At steady-state the concentration profile of the polarity cue along the cell is given by

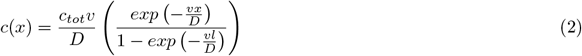

where *D* is the effective diffusion coefficient, and *c*_*tot*_ is the total amount of the conserved polarity cue (Fig.1D). For the full derivation of (Eq.2) see Section A in the Supplementary information (SI).

At the cell’s edges the polarity cue inhibits actin polymerization, and at steady-state the proportion of active polymerized actin is given by

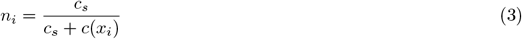

where 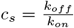 is the dissociation constant of the chemical reaction. For the full derivation of (Eq.3) see Section A in the SI. The polymerization speeds at both ends of the cell are written in the UCSP formulation

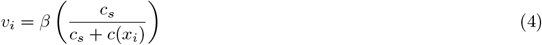

where *i* = *b, f* for either end of the cell. The net retrograde flow of actin for a cell of length *l* at steady-state (Fig.1C,D), where the boundaries are *x*_*b*_ = 0 and *x*_*f*_ = *l*, is therefore given by

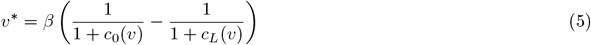

where *c*_0_(*v*), *c*_*L*_(*v*)=*c*(0)*/c*_*s*_, *c*(*l*)*/c*_*s*_ and we explicitly denote the dependence on the retrograde flow *v*. The dynamics of the net actin retrograde flow are given by the Langevin equation

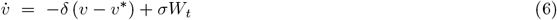

where *δ* is the rate at which the actin flow evolves to its steady-state value, *W*_*t*_ is a random Gaussian noise and *σ* is the noise amplitude. For the full derivation of this model see Section A in the SI.

We also consider that the speed of the cell is proportional to the retrograde flow (which was demonstrated for various type of cells in one-dimensional tracks [17, 27], and in keratocytes [28]), and in the opposite direction

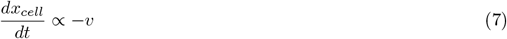

This follows from the observations that the traction force applied by the cell on the substrate is proportional to the retrograde flow of the actin [29]. In the over-damped regime of cells, this traction force is proportional to the resulting cell speed.

When studying the cell trains we treat cells as “glued” to each other, forming a single motile entity. The total traction force exerted by the train is given by the sum of all the traction forces of the individual cells, and the resulting speed is therefore proportional to this net force. To calculate the migration speed of a multi-cellular train (defined as the center-of-mass velocity *v*_*cm*_), we choose the simplest option where the speed is given by the average of the net retrograde flows of all the cells in the train.

Throughout this work the parameters of the model that we use were calibrated to fit the motion of isolated MDCK cells (Fig.1A), which migrate smoothly with (roughly) fixed length (for model calibration details see Section B, Fig.S1 in the SI).

## CELL DOUBLETS

### Introducing CIL and CL interactions between cells

Consider two UCSP cells with a fixed length *l*, which form an adhered pair which can not detach from one another. The net actin flow within each cell is denoted by *v*_*i*_ (*i* = 1, 2), and the local polymerization speed at the front/back of each cell *i* is denoted by *v*_*f/b,i*_. The local polymerization speeds, and the global retrograde flow within each cell depend on the concentration profile of the inhibitory cue *c*(*x*) (Eq.2).

The CIL and CL interactions of the pair are incorporated into the model by introducing two new parameters, which modify the inhibition level of actin polymerization at the edges of the cells that are in contact: 1) The parameter *a* for CIL, which symmetrically enhances the inhibition of actin polymerization activity at the touching cell edges *v*_*f*,1_ and *v*_*b*,2_ (Fig.2A) [5], and 2) The parameter *q* for CL, which enhances/reduces inhibition of actin polymerization at the touching edge with a lower/higher polymerization speed (Fig.2B). We assume here that the CL forms at the cell-cell contact by the cell edge that has locally higher actin polymerization activity. This cell therefore “wins” and is able to extend a CL, which was observed experimentally to exhibit enhanced and more sustained Rac1 activity, and hence higher actin polymerization [13]. The “losing” cell was observed to have its local actin polymerization activity diminished next to the CL of the “winning” cell [13, 16]. The interactions between the cells are mediated by various molecular components [3], which affect the actin polymerization activity on both sides of the cell-cell junction.

**FIG. 2:**
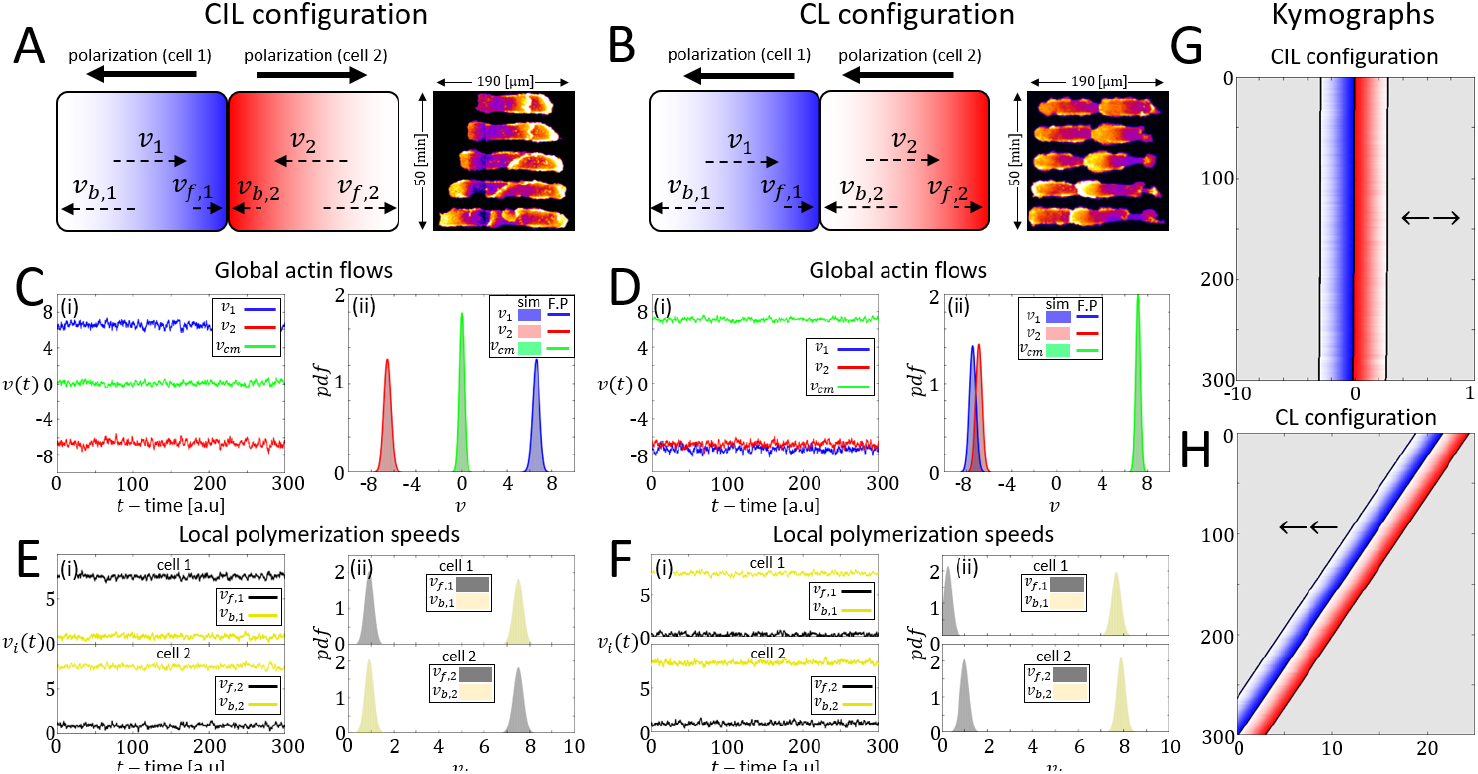
The cell pair model. A-B) Left panels: Illustration of the CIL and CL configurations in our model. Right panels: Short kymographs of MDCK cell expressing YFP-PBD signal while performing CIL and CL. Blue/Red colors indicate cell 1/2. *v*_1_/*v*_2_ are the speed of the actin retrograde flows (Eqs.8,9). *v*_*f*_ /*v*_*b*_ refer to the local polymerization speeds at the front and back respectively. Purple/Yellow fluorescence indicate a low/high intensity. C-D) The steady-state flows in the CIL configuration (*q* = 1 and *a* = 1) and the CL configuration (*q* = 4 and *a* = 0). (i) Time series of the global flows and the center-of-mass velocity (*v*_*cm*_). (ii) probability-density-functions for the global flows and the center-of-mass velocity. Blue/Red and green indicate the global flow in cell 1/cell 2 and the center-of-mass velocity. Solid curves in (ii) indicate the Fokker-Plank solutions. E-F) The local polymerization speeds in the CIL configuration (*q* = 1 and *a* = 1) and the CL configuration (*q* = 4 and *a* = 0). (i) Time series of the local actin polymerization speeds in cell 1 and cell 2. (ii) probability-density-functions for the local polymerization speeds in cell 1 and cell 2. Black/Yellow indicate the front/back. G-H) Simulated kymographs of a cell pair in a CIL and CL configuration respectively. Parameters: *l* = 2.8, *c* = 4, *D* = 4, *β* = 8, *δ* = 1, *σ* = 1*/*4.

Implementing these two interactions into the model, the steady-state global flows for two coupled cells are therefore given by (Eq.5)

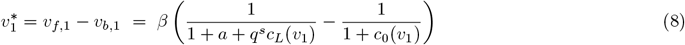

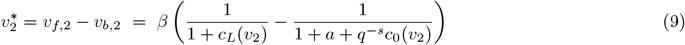

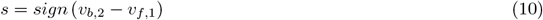

Note that we chose to implement the asymmetric CL interaction through a simple reciprocal relation between the effect of *q* on both edges (Eq.10). Throughout this work, the cell in which the inhibition of actin-polymerization is enhanced/reduced due to the effect of *q*, will be referred to as the losing/winning cell at the respective cell-cell contact. Our choice of implementing the CIL and CL interactions through this particular functional dependence on *a, q* should not affect the results in any qualitative way. Other functional forms that describe the symmetric inhibition for CIL, and asymmetric inhibition/enhancement for CL, should result in the same qualitative behavior.

We first examine the effects of these interactions, by studying the “pure” cases where only CIL or only CL affect the actin polymerization at the touching edges. The trajectories (Fig.2G) and the steady-state velocity distributions (Fig.2C,E) show that in the case of pure CIL the actin flows in both cells are opposite and symmetric and the cell pair remains stationary. For the case of pure CL the two cells are polarized and migrate in the same direction (Fig.2H), and the global actin flow in the losing cell (which is the leader cell of the pair) is faster (Fig.2D,F). In the CIL case the polymerization speeds at the touching edges of the cells are low and symmetric (Fig.2C), while in the CL case the polymerization speed in the losing edge is almost totally inhibited (Fig.2F). An analytic calculation of the velocity distributions, by solving the Fokker-Plank equations corresponding to Eqs.6,8-10, is shown in SI C. In Fig.2A,B we display short experimental kymographs of MDCK cell pairs exhibiting CIL and CL configurations respectively.

### CIL-CL phase diagram

Next, we span the cell doublet *a* − *q* phase diagram (Fig.3A), by analyzing the steady-state solutions of the deterministic equations for the global flows *v*_*i*_ of the individual cells in the cell pair (Fig.3B, Eqs.8-10). The *a* – *q* phase-diagram depicts the possible steady-state configurations which cell doublets can assume with respect to different sets of *a* and *q*. For more analysis of (Eqs.8-10) see Section D in the SI.

**FIG. 3:**
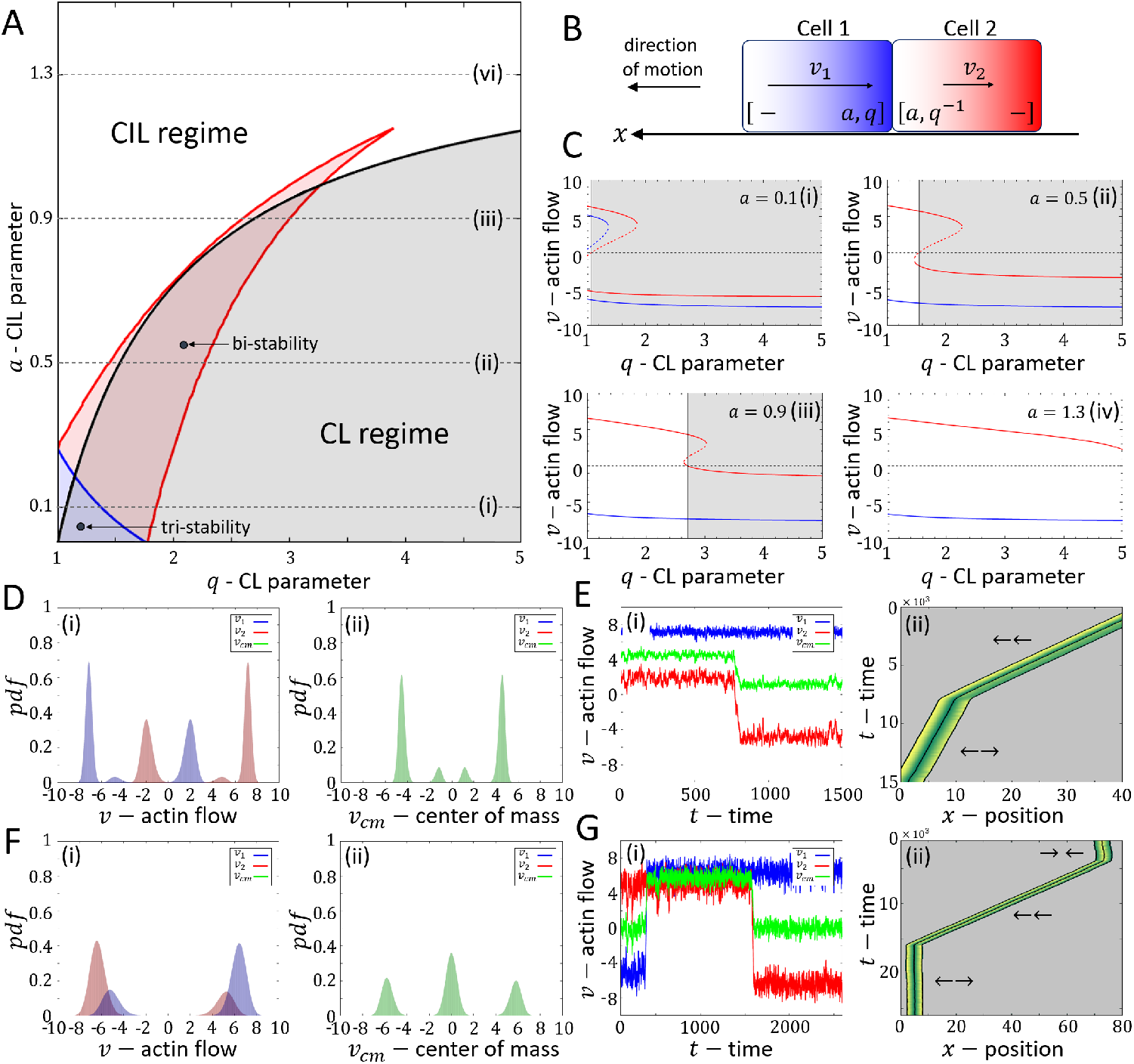
A) The *a* − *q* phase-diagram for cell pairs. The diagram assumes a CL configuration in which cell 2 is the winning cell, as depicted in (B). Blue/Red curves indicate saddle node bifurcations for cell 1/cell 2 respectively. Black curve indicates the *a*_*c*_ transition line (Eq.11) where the global flow in cell 2 changes a direction (red cell in B). B) Illustration of a polarized CL configuration of a cell pair. Solid arrows indicate the direction of the global internal flows in each cell. C) panels (i-vi) correspond to the steady-state solutions (Eqs.8-10) of *v*_1_ (blue) and *v*_2_ (red) as a function of *q* for the values of *a* = 0.1, 0.5, 0.9, 1.3 indicated on the phase-diagram (black dashed lines in A). D) probability-density-functions for (i) the global flows *v*_1_*/v*_2_ in blue/red and (ii) the center-of-mass velocity *v*_*cm*_ in green in the bi-stable region ((*a, q*) = (0.6, 2.1), black point in A). Noise amplitude is *σ* = 1*/*4. E) Example of a time series for the bi-stable point (i) and its corresponding kymograph (ii). Blue/Red indicate the global flows *v*_1_/*v*_2_ and green indicate the center-of-mass velocity *v*_*cm*_. F) probability-density-functions for (i) the global flows *v*_1_*/v*_2_ in blue/red and (ii) the center-of-mass velocity *v*_*cm*_ in green in the tri-stable region ((*a, q*) = (0.08, 1.12), black point at in A). Noise amplitude is 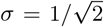. G) Example of a time series at the tri-stable point (i) and its corresponding kymograph (ii). Blue/Red indicate the global flows *v*_1_/*v*_2_ and green indicate the center-of-mass velocity *v*_*cm*_. Parameters: *l* = 2.8,*c* = 4,*D* = 4,*δ* = 1.

In the *a* − *q* phase diagram we find that for large values of *a*, the stable configuration is CIL (←→), as the two steady-state solution branches of *v*_1_ and *v*_2_ (Eqs.8-10) oppose one another in sign, for all values of *q* (Fig.3C,iv). As *a* decreases, we find two transition lines which correspond to a fold that appears in the solution branch of the winning cell (at the rear, Fig.3B) *v*_2_ (red curves in Fig.3C,i-iii). This fold in the solutions creates a region of bi-stability between the CIL and CL (→→) states (region marked by the red curves in Fig.3A). Following a further decrease in *a*, or alternatively an increase in *q*, beyond the fold in the solution curve, the stable configuration is only CL (region of *q* where the blue and red curves are of the same sign in Fig.3C,i-iii). For very low values of *a* and *q* we find another transition line which is associated with the solution branch of the losing cell *v*_1_ (blue transition line in Fig.3A). This transition line denotes a saddle-node bifurcation (Fig.3C,i), and below this transition line the effects of the interaction parameters are weak and the system can assume all the CIL, CL, and anti-CIL (a-CIL, →←) steady-state configurations.

We also find a fourth transition line, which is associated with the change of the direction in the global flow inside the wining cell *v*_2_ (black curve in Fig.3A). The change in flow direction corresponds to the limit of *v*_2_ *→* 0 (marked by the transition from white to grey shade in Figs.3C,ii-iii), and this transition line is therefore given by the following approximate relation for the critical values of *a* and *q*

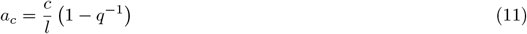

We find that this transition line is highly significant, as above this transition line (*a* > *a*_*c*_), all the configurations are CIL. In the narrow regions of bi-stability above this transition line the cell pair can assume two different CIL states, where the cell pair is nevertheless motile and persistent (red curve in Fig.3C,(iii)). For the derivation of (Eq.11) see see Section D in the SI.

The richness of the migration patterns of the cell pair in our model is demonstrated when we examine the regions of bi-stability and tri-stability (at the black points in Fig.3A). In the bi-stable region (red region in Fig.3A)), the steady-state velocity distributions (Fig.3D,i-ii), the internal flow trajectories and their corresponding kymograph (Fig.3E,i-ii) demonstrate that there are two distinct persistent motility phases, one is in a CL configuration and the other, with a slower pair velocity, is in a CIL configuration. In the tri-stable region, where the interactions are weak (blue region in Fig.3A), the steady-state velocity distributions (Fig.3F,i-ii), the internal flow trajectories and corresponding kymograph (Fig.3G,i-ii) demonstrate that there is a motile CL configuration, along with two distinct non-motile phases: one phase is in a CIL configuration and the other is an a-CIL configuration. We note that as long as there is some CL interaction (*q* > 1), the CIL configuration will have some residual speed and the pair will be motile. An extended analysis of the doublet dynamical behavior across the *a* − *q* phase-diagram is presented in Section E in the SI.

### Effects of noise and comparing to experiments

In order to understand the role of fluctuations (noise) in the polymerization speeds, we study the steady-state behavior for: 1) a low value of noise, which is close to the deterministic solutions which we described in Fig.3 (Fig.4A), and 2) a high value of noise, which may exist in cells (Fig.4B). For this purpose, we plot for two level of noise, the configuration probability heatmaps for each configuration that the doublet system can assume: CL (Fig.4A,B,i), CIL (Fig.4A,B,ii), and a-CIL (insets in Fig.4A,B,ii), as well as the pair’s mean center-of-mass velocity heatmaps (Fig.4A,B,iii). For the computational method used to calculate the proportions of the configurations, and the proportions of internal states within a configuration from the simulations, see Section G in the SI.

**FIG. 4:**
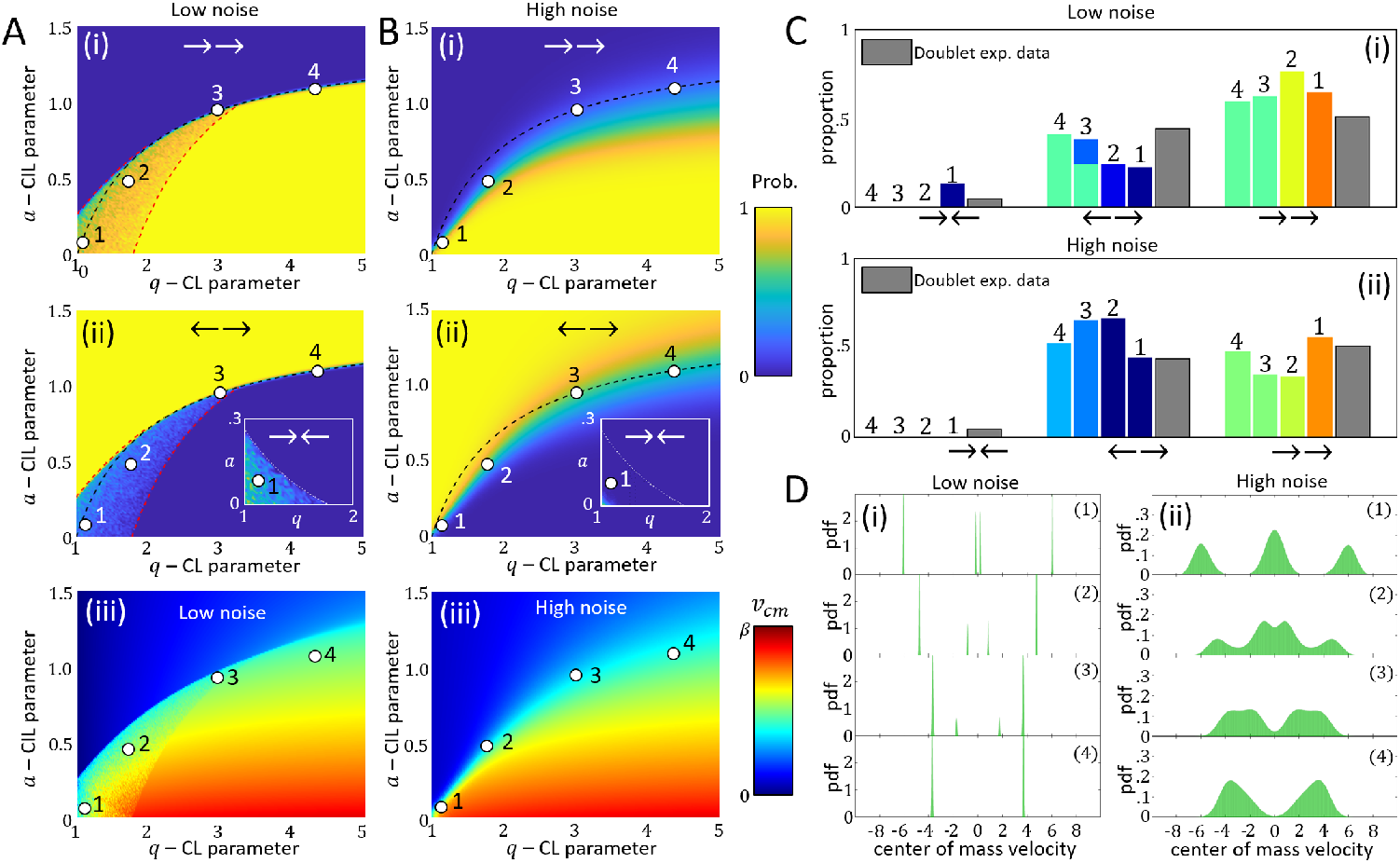
A-B) Low and High noise analysis. (i) CL configuration probability heatmap. (ii) CIL configuration probability heatmap. right-low inset is the a-CIL probability configuration heatmap. Yellow/Blue colormap indicates a probability 1/0 (iii) center-of-mass velocity heatmap. Red/Blue colors indicate *v*_*cm*_ = *β*/*v*_*cm*_ = 0. In (A) black, red and white dashed curves are the analytical bifurcation transition lines shown in Fig.3A. In (B) the dashed black line denotes the analytic *a*_*c*_ transition line (Eq.11). Red/Blue colormap indicate *v*_*cm*_ = *β*/*v*_*cm*_ = 0. C) Configuration probability for low (i) and high (ii) noise amplitudes with respect to the points labeled by 1-4 in (A-B). Gray color indicate the experimental data [16]. Other colors indicate the center-of-mass velocity with respect to the color-map in (A-B,iii). D) Center-of-mass velocity probability-density-functions for low (i) and high (ii) noise amplitudes with respect to points 1-4 in A,B. Low and high noise amplitude are taken to be *σ* = 1*/*32 and 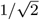 respectively. Other parameters: *l* = 2.8, *c* = 4, *D* = 4, *β* = 8, *δ* = 1. Point 1-4 correspond to (*a, q*) = (0.08, 1.12), (0.48, 1.74), (0.95, 3), (1.1, 4.33).

The probability heat-maps show that for a low level of noise, the steady-state dynamics of the cell pair corresponds to the deterministic transition lines, as predicted in the *a − q* phase-diagram (Fig.3A), whereby the bi-stable region between the CIL and CL phases can be identified (Fig.4A,i-ii), as well as the tri-stable regime which includes the a-CIL configuration (inset in Fig.4A,ii). For a high level of noise the heat-maps simplify into CIL-dominated and CL-dominated regions, separated approximately by the *a*_*c*_ transition line (Eq.11) (Fig.4B,i-ii). These findings also correspond to the mean center-of-mass velocity heat-maps, which shows a transition from low to high pair migration velocities in the proximity of the *a*_*c*_ transition line (Fig.4A,B,iii).

We next compare between the probability distribution of the different polarization configurations of MDCK cell doublets (measured by the polarity signal YFP-PBD, Fig.2A,B), and our model simulations in the proximity of the *a*_*c*_ transition line (points 1-4 in 4A-B). The internal polarization of the coupled MDCK cells are determined by the difference in intensity of the PBF-YFP signal at the cell boundaries (see see Section F in the SI for experimental methods). The experimental data [16] (Gray bars in Fig.4C) shows that the CIL and CL configurations have similar proportions, while the transient a-CIL configuration has very small proportions. The simulations demonstrate that along the *a*_*c*_ transition line the CIL and CL configurations have relatively comparable weights, similar to the experiments. The transient a-CIL phase however, only appears for simulations with low values of noise in the multi-stable point (point 1 in Fig.4C,i), while at a high level of noise this transient phase is unstable. The simulation results also highlight that in the bi-stable region we find two CIL phases, with different speeds of the pair’s center-of-mass (point 3 in Fig.4C,i). The center-of-mass velocity distributions (Fig.4D) demonstrate how the transient a-CIL phase of low velocity magnitude decreases the pair’s migration in the CIL phase when the noise is increased. Despite to the limited data set, and the number of free parameters in the mode, our analysis suggests the following qualitative conclusion, that the MDCK cells are poised close to the CIL-CL transition line.

## CELL TRIPLETS

### The middle-cell and the triplet phase-diagram

Next, we expand the model to describe cell triplet trains. In the triplet system, we have a new type of “bulk”, middle cell (cell 2), which is “inserted” between the two edge cells. The cells at the edges (now labeled as ‘cell 1’ and ‘cell 3’, Fig.5A bottom panel) are described by the same polarization states as in the doublet. The middle cell can assume polarized states, as well as two non-polarized states where both edges are either inhibited or activated symmetrically, due to its interactions with the neighboring cells.

**FIG. 5:**
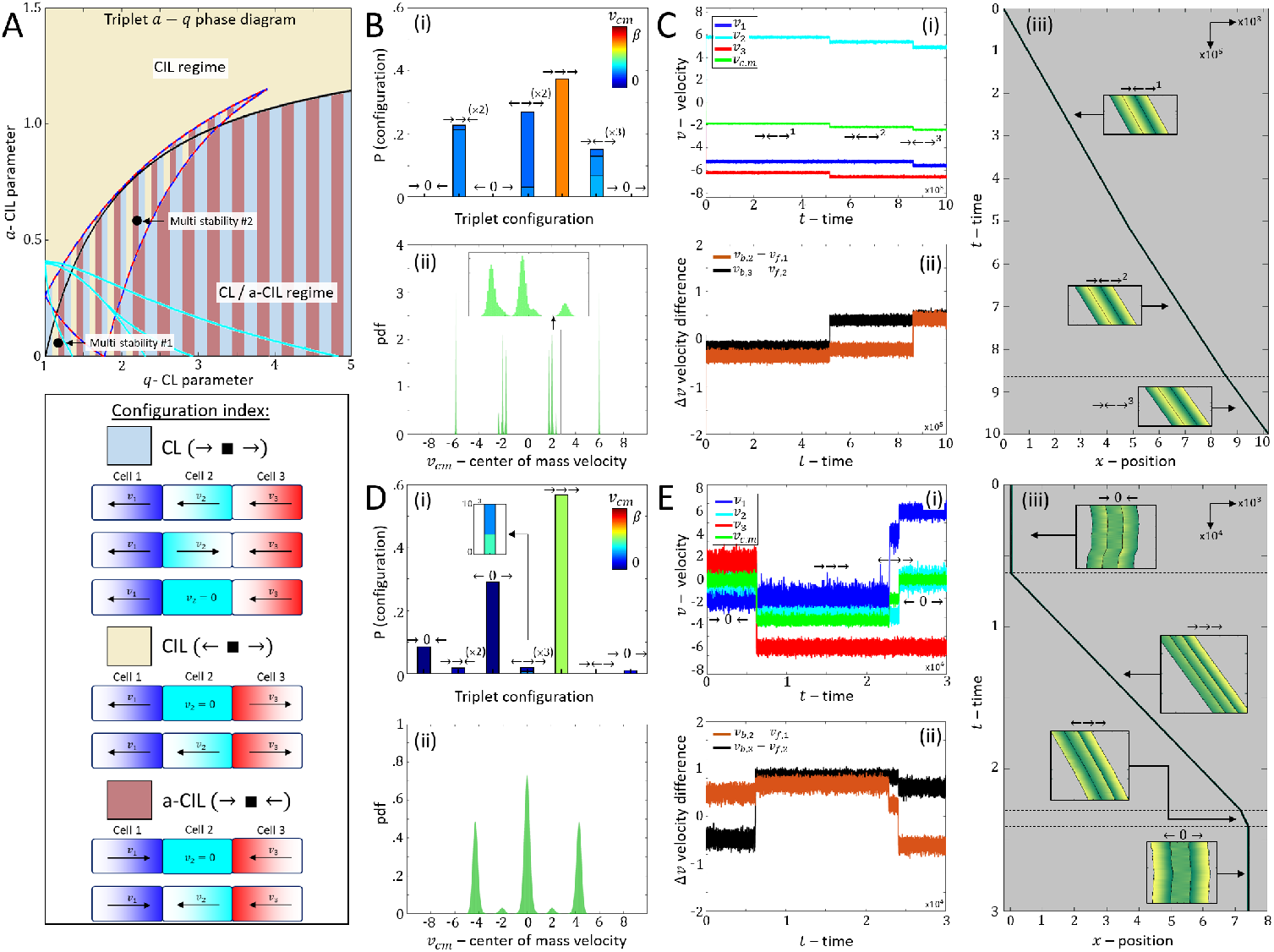
A) The triplet *a*-*q* phase-diagram. The diagram accounts a configuration of three cells over which cell 1 (blue) and cell 3 (red) are the edge cells and their analysis is similar to the cell doublets (as in Fig.3A-B without a pre-determined CL configuration), and cell 2 (teal) is the middle cell which can assume three independent internal configurations (see Section H in the SI). Striped blue-red curves and teal curves indicate the saddle node bifurcation transition lines for the cell 1-cell 3 and cell 2 respectively. Black dashed curve indicates the *a*_*c*_ transition line (Eq.11) over which the global flow in the edge cell over which the actin polymerization speed is enhanced changes a direction (either cell 1 or cell 3). Khaki/light blue/brown represent regions of CL/CIL/a-CIL configurations (defined by the edge cells, as depicted in the configuration index below). Black-white points indicate (*a, q*) = (0.08, 1.12) and (*a, q*) = (0.6, 2.1). B) steady-state behavior for (*a, q*) = (0.08, 1.12). (i) The triplet configuration probability. Color code indicates the mean center-of-mass velocity *v*_*cm*_. (ii) probability-density-function of the center-of-mass velocity *v*_*cm*_ in green. C) Multi-stable trajectory example from point 2 ((*a, q*) = (0.08, 1.12)). (i) Time series of the global flows, which demonstrates a case where the train transitions between different internal flows within the configuration → ← →. Blue/Teal/Red is the global flow in cell 1/2/3. Green is the center-of-mass velocity. (ii) The difference between the local polymerization speeds at the touching edges. Black/Orange stand for the contact between cells 2-3/ cells 1-2 respectively. (iii) Corresponding kymograph indicating the transitions between the internal flows. D) steady-state behavior for (*a, q*) = (0.60, 2.10). (i) The triplet configuration probability. Color code indicates the mean center-of-mass velocity *v*_*cm*_. (ii) probability-density-function of the center-of-mass velocity *v*_*cm*_ in green. E) Multi-stable trajectory example from point 1 ((*a, q*) = (0.08, 1.12)). (i) Time series of the global flows, which demonstrates a case where the train transitions between CIL (← 0 →) and CL (→ → →) states through a transient ← → → state. Blue/Teal/Red is the global flow in cell 1/2/3. Green is the center-of-mass velocity. (ii) The difference between the local polymerization speeds at the touching edges. Black/Orange stand for the contact between cells 2-3/ cells 1-2 respectively. (iii) Corresponding kymograph indicating the transitions between the internal flows. Parameters: *l* = 2.8, *β* = 8, *c* = 4, *d* = 4, *δ* = 1, Noise levels: *σ* = 1*/*8 in C, *σ* = 1*/*4 in F. Yellow/Green regions in kymographs indicate high/low inhibitory cue concentration.

The self-consistent equations for the steady-state flows inside the cells of the triplet, are given by

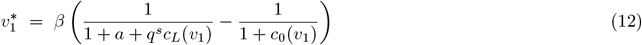

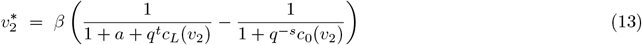

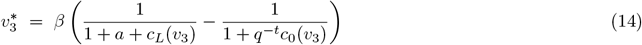

where *t, s* = ±1.

For this system the *a*−*q* phase-diagram is obtained by solving (numerically) the deterministic steady-state solutions of the global actin flows (Eqs.12-14), when considering all the possible configurations for the interactions within the triplet (Fig.5A). This process is done by constructing separate phase-diagrams for each internal configuration of the middle cell, and is fully presented in Section H in the SI.

We choose to classify the configurations with respect to the behavior of the cells at the edges of the train (Fig.5A bottom panel): 1) CL (→ □ →), 2) CIL (← □ →), and 3) a-CIL (→ □ ←). Within this classification the middle cell, denoted by □, can assume either a polarized state (→,←) or a non-polarized state (0). After considering all the symmetries in the system, we find that the triplet system can assume seven configurations (Fig.5A).

We find that similar to the doublets phase-diagram (Fig.3A), the CIL and CL configurations in the triplet *a – q* phase-diagram are mostly divided by the *a*_*c*_ transition line (black curve in Fig.5A). Above this transition line (large *a*) all the configurations that the triplet can assume are CIL (khaki color in Fig.5A), while below the *a*_*c*_ transition line (low *a* large *q*) the CL and a-CIL configurations span the configuration space (Light blue and Brown regions in Fig.5A). The triplet phase diagram also displays new transition lines due to the multiple solution that the middle cell (*v*_2_) can assume (teal curves in Fig.5A), as well as the transition lines due to the solutions of the edge cells (*v*_1_ and *v*_3_, red-blue striped curves in Fig.5A). In between these transition lines we find that the system is multi-stable and can assume many configurations with different probabilities (see Section I in the SI for the full analysis of the triplet *a − q* phase diagram). The stability of each configuration is reflected in the proportion of time the system spends in this state, as discussed below.

To demonstrate the rich variety of triplet dynamical behaviours in the multi-stable regimes, we explore using numerical simulations the same regions which were studied for the doublet case (black points in Fig.5A for the triplets, and Fig.3A for the doublets). For the region of weak interactions (low *a* and *q*) we find that the system has several stable solutions for each polarization configuration (Fig.5B): we demonstrate a trajectory where the triplet train remains in the same polarization configuration (→←→), while it switches between three different internal states with slightly different migration speeds (Fig.5C). See Section G in the SI for the procedure of calculating the proportions of the internal states from the model simulations.

In the second region of multi-stability (where the interactions are stronger), we also find that the system has several stable solutions for each polarization configuration (Fig.5D). We demonstrate how a triplet transitions between motile and stationary configurations along a single trajectory (Fig.5E), where the system transitions between a-CIL (→ 0 ←), to full CL (→→→), and then to full CIL (← 0 →) configuration through a transient CIL configuration (←→→). For both of these examples we demonstrate that the transitions between the motility modes occur due to changes in the local polymerization speeds at the touching edges of the cells (Fig.5C,E,ii). An extended analysis of the steady-state behavior along other multi-stable regions in the triplet *a*-*q* phase-diagram is presented in Section I in the SI.

### Effects of noise and comparison with experiments

Next, we plot the configuration distribution heat-maps for low and high levels of noise amplitudes on the *a − q* phase diagram (Fig.6A-C,F-H), as well as the mean center-of-mass velocity heatmaps (Fig.6,D,I). The heatmaps are depicted for the individual configurations (panels (i-ii) in Fig.6A-C,F-H), and when classified according to the edge cells’ polarization (panels (iii) in Fig.6A-C,F-H). For a low level of noise we find that, similar to the doublets, the deterministic transition lines of the phase-diagram (Fig.5A) are captured in the heatmaps (Fig.6A-C). For a high level of noise we find that the transient configurations disappear and the heatmaps simplify into two main regions that are divided in the proximity of the *a*_*c*_ transition line. For large values of *a* the heatmaps are dominated by CIL, while for low *a* and large *q* (below the transition line) the heatmaps are dominated by the CL configuration. We also find that for large values of *q* the proportions of the a-CIL configuration increase as the a-CIL solution is stabilized in this regime (see Section H in the SI for analytical explanation). The appearance of the a-CIL modes for large values of *q* also affects the center-of-mass velocity heatmaps. In the CL regime (low *a*) the triplet average speed peaks (red regions in Fig.6D,I), and then decreases with increasing *q* (Fig.6D,I), due to the increased likelihood of weakly motile a-CIL configurations (Fig.6C,H).

**FIG. 6:**
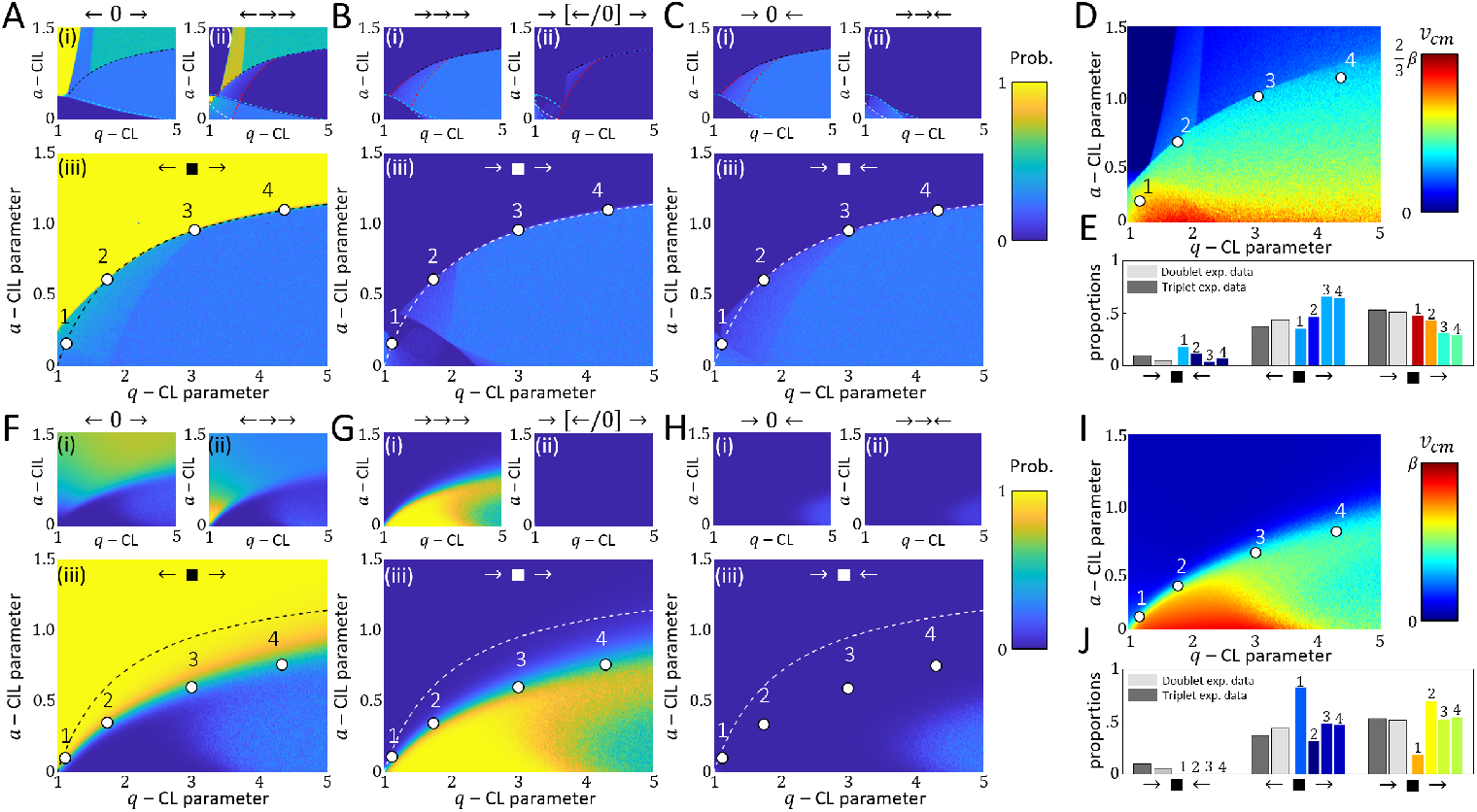
Triplet noise analysis. A-D) Low noise (*σ* = 1*/*32): A) *a* − *q* CIL configuration heatmaps where: (i) the middle cell is not polarized, (ii) the middle cell is polarized, and (iii) all the configurations in which the edge cells act as CIL. Vertical lines in the upper-left are a numerical artifact due to the velocity threshold which was used to separate between a finite and zero velocities of the middle cell. B) *a* − *q* CL configuration heatmaps where (i) the middle cell is polarized in the same direction as the edge cells, (ii) the middle cell is not polarized or polarized against the direction of the edge cells, and (iii) all the configurations in which the edge cell act as CL. C) *a* − *q* a-CIL configuration heatmaps where (i) the middle cell is not polarized, (ii) the middle cell is polarized, and (iii) All the configurations in which the edge cells act as a-CIL. D) *a* − *q* center-of-mass velocity heatmap. E) The configuration distribution for points 1-4 indicated in D compared to the experimental data of the doublets and the triplets. Colors represent the center-of-mass velocity with respect to the color code in D. Light/Dark gray bars indicate the experimental data for the doublets/triplets. F-J) High noise 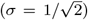: Same as panel description as in A-E above. Parameters: *l* = 2.8, *β* = 8, *c* = 4,*d* = 4, *δ* = 1. Points 1-4 in A-D: (*a, q*) = (0.15, 1.12), (0.60, 1.74), (0.95, 3.00), (1.09, 4.33). Points 1-4 in E-H: (*a, q*) = (0.1, 1.12), (0.34, 1.74), (0.6, 3), (1, 4.33).

In our model, the a-CIL configuration is stabilized at large *q* values since the cryptic lamellipodia are enhanced by, and maintain, the non-polar state of the middle cell. There are experimental indications that cells can indeed maintain a persistent polarity while pushing against their soft neighbors [13, 16] or soft barriers [30]. However, there may also be other mechanisms or signals that make it easier for the edge cells to protrude outwards, or that destabilize the non-polar state of the middle cell (converting it to a polarized cell), thereby limiting the stability and frequency of the a-CIL configuration in experiments.

We compare between the experimental observation of the proportions of triplet polarization states, collected for cell triplets using the PBD-YFP intensity measurement (see Section F in the SI for experimental methods), and the simulation results along the *a*_*c*_ transition line (Fig.6E,J). Note that for the simulations with a high level of noise, the data was collected below the *a*_*c*_ transition line, as the heatmaps show that the actual transition between the CIL and CL regions is shifted to lower values of *a* (Fig.6F-H,ii) in comparison to the doublets and triplets of low noise (Figs.4B,6A-C). For the triplet configurations that are classified by the edge cells’ polarization, the proportions of the cell doublets and triplets distribute similarly (light and dark gray bars in Fig.6E,J). We demonstrate that, as in the case of the doublets ((Figs.4C), in the proximity of the *a*_*c*_ transition line the simulated proportions of the configurations of the triplets are similar to the experiments (points 1-4 in Fig.6E,J). These results suggests, as in the case of doublets, that cells are poised close to the critical transition line where CIL and CL interactions are balanced.

## COLLECTIVE MIGRATION OF LONG CELL TRAINS

We now explore the migration modes of longer cellular trains composed of *N* cells (Fig.7A). To construct longer trains in the model, we insert cells in the middle of the train (the train’s bulk) which can assume the same internal configuration as the middle cell in the triplet train (Eq.13), and therefore there is no new *a − q* phase diagram to depict, as there are no new functionals to explore, beyond those in Eqs. 12-14.

**FIG. 7:**
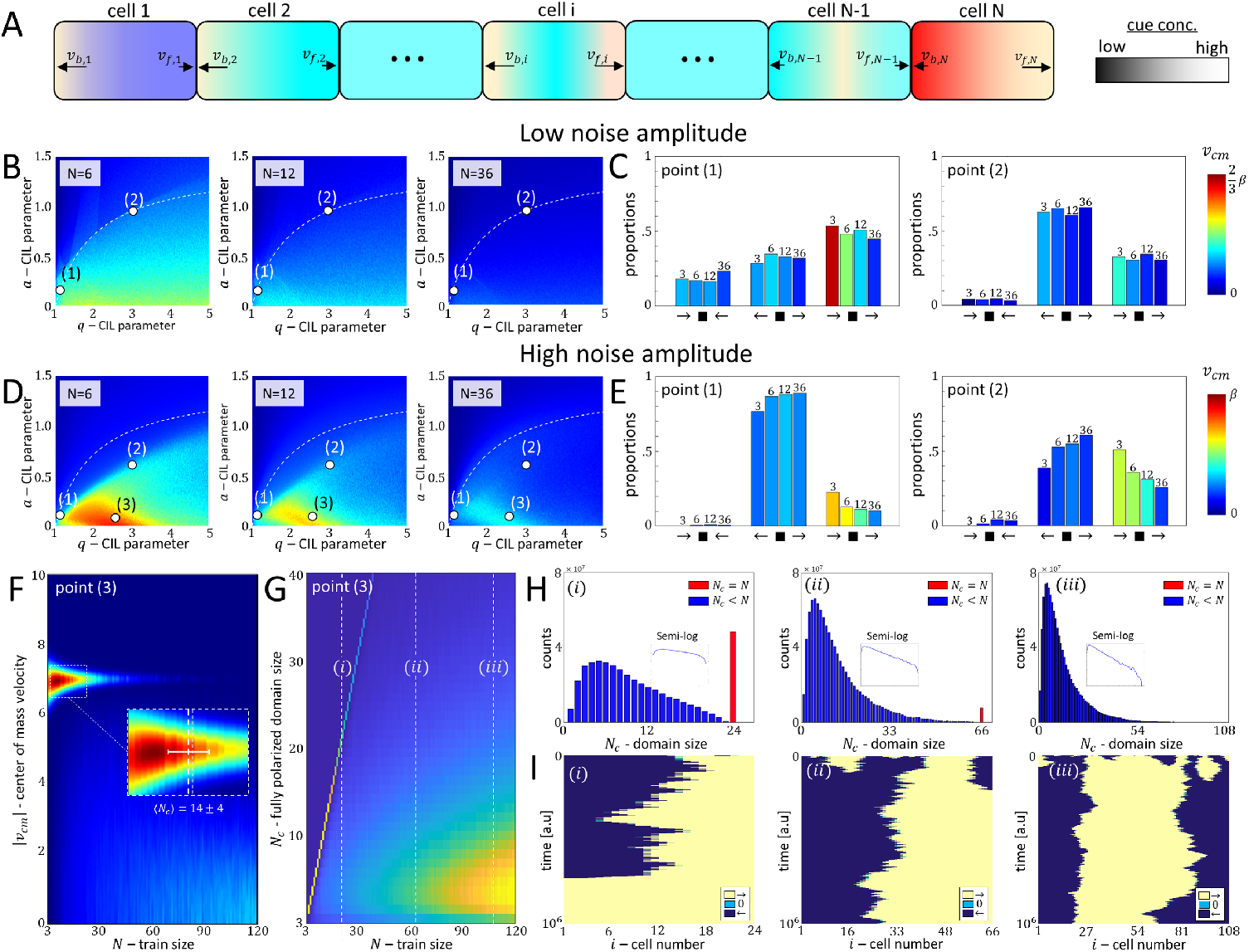
Analysis for long trains of *N* size. A) Illustration of a cell train of size *N*. Blue/Red indicate the edge cells (cell 1 and cell N). Teal indicate the cells in the train’s bulk. Color gradient indicates the inhibitory cue concentration profile. B) Mean center-of-mass velocity heatmaps for an increasing train size (*N* = 6, 12, 36) as a function of *a* and *q* for a low noise level (*σ* = 1*/*32). White point (1)/(2) indicates (*a, q*) = (0.153, 1.12)*/*(0.95, 3). C) The configuration proportions for a system size of *N* = 3, 6, 12, 36 at points 1-2 indicated in (B). Colors in B and C represent the mean center-of-mass velocity with respect to the color code on the right. D) Mean center-of-mass velocity heatmaps for an increasing train size (*N* = 6, 12, 36) as a function of *a* and *q* for a low noise level 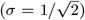. White Point (1)/(2)/(3) indicates (*a, q*) = (0.1, 1.12)*/*(0.6, 3)*/*(0.05, 2.5). E) The configurations proportion for a system size of *N* = 3, 6, 12, 36 at points 1-2 indicated in (D). Colors in D and E represent the center-of-mass velocity with respect to the color code on the right. F) Mean center-of-mass velocity distribution heatmap (in absolute value) as a function of train size *N* in point (3) in (D). Red/Blue color gradient indicate regions of high/low density. G) Fully polarized domain size (*N*_*c*_) distribution heatmap as a function of system size *N* for point (3) in (D). White dashed horizontal lines indicate a system size of *N* = 24, 66, 108. Yellow/Blue color gradient indicate regions of high/low density. H) Domain size distribution for trains of size *N* = 24*/*66*/*108 in (i)/(ii)/(iii) respectively (White dashed lines in B). Red/Blue bars indicate the distributions of domains which are equal to/smaller than the system size. Small insets display the semi-log plot of the cluster domain distribution excluding domains of system size (blue bars only). Linear fitting parameters for the inverse of the slope for (i)/(ii)/(iii) is: 12.01*/*10.75*/*9.17 respectively, corresponding to the typical correlated cluster size. I) Configuration dynamics for cell trains. (i)/(ii)/(iii) indicate *N* = 24*/*66*/*108 respectively (white dashed lines in E). Dark Blue/Bright Yellow/Teal colors indicate cells with positive/negative/zero polarization. Parameters: *l* = 2.8,*d* = 4,*c* = 4.

We explore using numerical simulations the steady-state behavior of trains of *N* cells, for both small and large values of noise in the actin polymerization speed by calculating the center-of-mass velocity heatmaps for low and high values of noise, and calculate the proportions of the configurations by classifying the configurations by the polarization of the edge cells (as in the case of the triplets). We depict the mean center-of-mass velocity heat-maps for increasing train sizes (Fig.7B), which for low noise demonstrate the same trend observed in the doublets and triplets: The CIL and CL configurations are divided near the *a*_*c*_ transition line, independent of the train size. For high noise, the CIL-CL transition is shifted to lower values of *a* (Fig.7D). We also find that similar to the triplets, the train speed peaks at some intermediate *q* value and decreases for large values of *q* (Fig.7B,D). This occurs due to the increase in the proportion of the a-CIL configurations as *q* increases (Fig.7C,E). We find that for low noise the proportions of the polarization states of the trains do not change significantly with increasing train length (Fig.7C), at higher noise there is a clear trend of longer trains becoming less polarized and less motile, with higher CIL and a-CIL proportions (Fig.7E).

The region which is identified with the most polarized CL configuration, highest *v*_*cm*_ in Fig.7(D), diminishes as the train size grows. In this regime (marked by a white dot in Fig.7(D), *N* = 6) we plot a heatmap of the center-of-mass velocity as a function of system size (Fig.7F). We find that the highest velocity correlates with train size of *N ≈* 3 – 20 (inset in Fig.7F), and as the train size grows further, the mean velocity distribution shifts to lower velocities. The origin of these changes is explained by plotting the distribution of the sizes of polarized cell clusters *N*_*c*_ (the number of neighbouring cells which are polarized in the same direction) as a function of system size (Fig.7G), and the cluster size distributions (Fig.7H) for the sizes indicated by the white dashed lines in Fig.7G. In the distributions of the cluster size we note the case where the whole train forms a single polarized domain (*N*_*c*_ = *N*, red bars in Fig.7H), and when clusters of polarized domains are smaller than the system size (*N*_*c*_ < *N*, blue bars in Fig.7H).

The results show that as the train size increases, the probability of a train to be in a single polarized domain diminishes (Fig.7G,H), while the distribution of the cluster domains of smaller size *N*_*c*_ < *N* increases in its weight. This distribution obeys an exponential (Poisson) distribution (insets in Fig.7H), implying that the domain walls are independent and have a finite probability to be randomly generated along the train. The persistent cluster size analysis reveals that the mean persistent cluster size is *N* = 14 ± 4, which correlates region of high velocity (inset in Fig.7F). This is similar to the typical size of the correlated cluster size obtained from the cluster size distribution (Fig.7H) In Fig.7(I) we demonstrate typical cluster dynamics of trains of various sizes (indicated by the white dashed lines in Fig.7G). The simulations show that the cells within the domains are fully polarized, and non-polarized cells appear as single cells domain walls between the domains of opposite polarization (Fig.7I).

In the large *q* regime, we find that the speed of long trains decreases (Fig.7B,D), similar to the triplets (Fig.6). The reason for this behavior is due to the proliferation of a-CIL domain walls, which break the train into numerous short domains, resulting in small overall coherence and global migration speed (see Section J in the SI).

In our model the persistence of the individual cell is determined by the UCSP parameters that determine the strength of the spontaneous cellular polarization [19]. We therefore expect that more persistent cells will give rise to correlated clusters of larger mean size.

## CELLULAR RINGS (TRAINS WITH PERIODIC BOUNDARY CONDITIONS)

In [13] it was observed that one-dimensional cellular trains switch from a less correlated to a more correlated collective motion when they form closed loops on ring-like tracks. We are therefore motivated to analyze the effect of periodic boundary conditions on our model system (Fig.8). For the full configuration distributions see Section J in the SI.

**FIG. 8:**
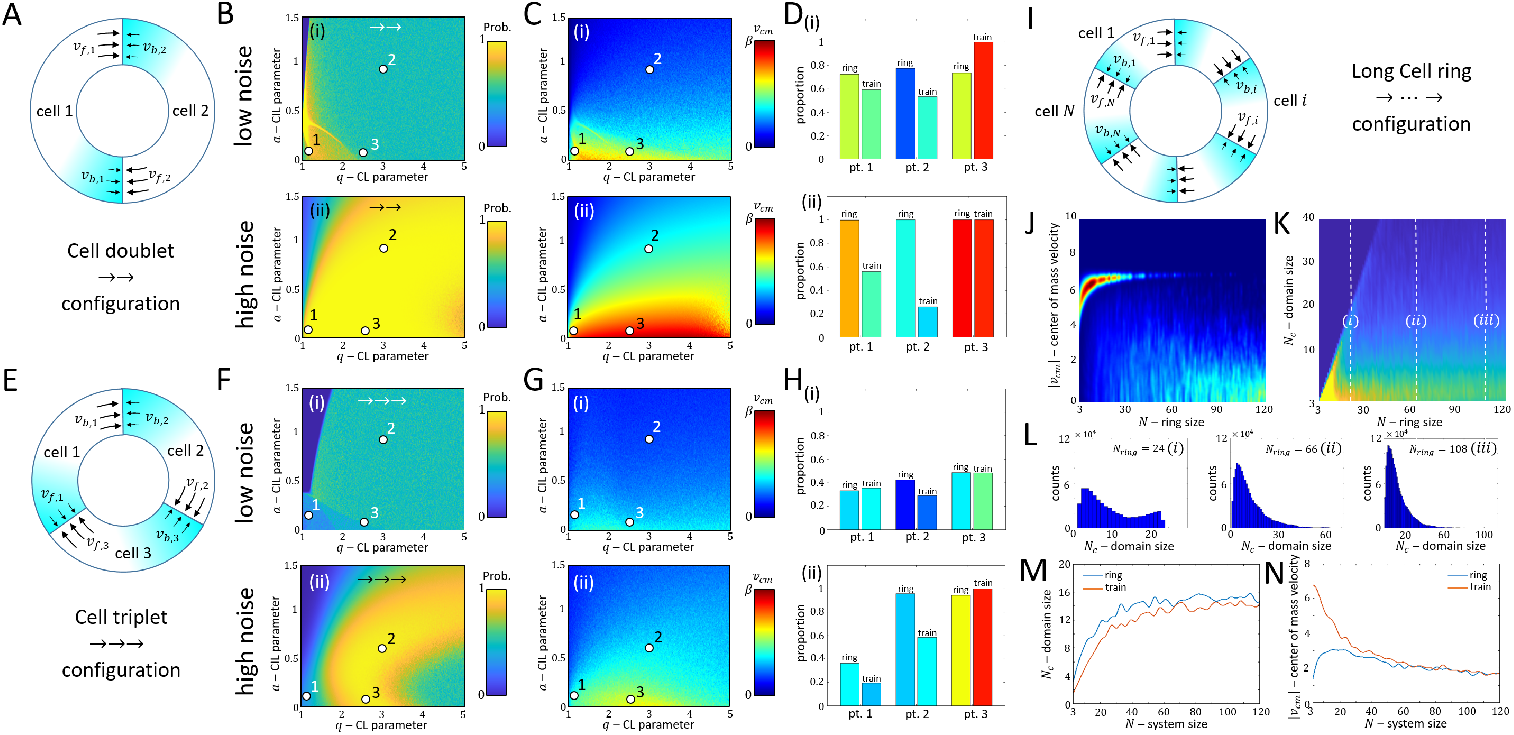
Model for *N* cells rotating on a ring. Cell doublet: A) Illustration of a persistent cell doublet rotating along a ring with a CL configuration. Color gradient indicates the inhibitory cue concentration profile. B) The → → configuration proportion distribution *a* − *q* heatmap for a low/high noise level (i)/(ii). Yellow/Blue colors indicates regions of high/low probability. C) The *a* − *q* center-of-mass velocity heatmap for a low/high noise level (i)/(ii). Color gradient indicates the magnitude of the center-of-mass velocity. Points 1/2/3 in (i)/(ii) refer to (*a, q*) = (0.08, 1.12)*/*(0.95, 3)*/*(0.05, 2.5). D) Comparison between the proportions of the → → configuration in a ring (periodic boundary conditions), and a train (free boundary conditions) for points 1,2,3 indicated in B and C for a low/high noise level (i)/(ii). Colors represent the mean center-of-mass velocity with respect to the color code to the left. Cell triplet: E) Illustration of a persistent cell triplet rotating along a ring with a CL configuration. Color gradient indicates the inhibitory cue concentration profile. F) The → → → configuration proportion distribution *a* − *q* heatmap for a low/high noise level (i)/(ii). Yellow/Blue colors indicates regions of high/low probability. G) The *a* − *q* center-of-mass velocity heatmap for a low/high noise level (i)/(ii). Color gradient indicates the magnitude of the center-of-mass velocity. Points 1/2/3 in F-G,(i) refer to (*a, q*) = (0.15, 1.12)*/*(0.95, 3.00)*/*(0.05, 2.5). Points 1/2/3 in F-G,(ii) refer to (*a, q*) = (0.1, 1.12)*/*(0.6, 3)*/*(0.05, 2.5). H) Comparison between the proportions of the → → → configuration in a ring (periodic boundary conditions), and a train (free boundary conditions) for points 1,2,3 indicated in B and C for a low/high noise level (i)/(ii). Colors represent the mean center-of-mass velocity with respect to the color code to the left. I) Illustration of a persistent cell train of *N* size rotating along a ring with a CL configuration. Color gradient indicates the inhibitory cue concentration profile. J) Mean center-of-mass velocity distribution heatmap (in absolute value) as a function of train size *N*. Red/Blue color gradient indicate regions of high/low density. K) Cluster size *N*_*c*_ distribution heatmap as a function of system size *N*. White dashed horizontal lines indicate a system size of *N* = 24, 66, 108. Yellow/Blue color gradient indicate regions of high/low density. L) Cluster size distribution for trains of size *N* = 24*/*66*/*108 in (i)/(ii)/(iii) respectively (White dashed lines in K). M) Mean center-of-mass velocity *v*_*cm*_ as a function of system size *N* for cell rings (blue) and cell trains (orange). N) Persistent domain size *N*_*c*_ as a function of system size *N* for cell rings (blue) and cell trains (orange). Parameters: *l* = 2.8,*d* = 4,*c* = 4.

We start with the analysis of cell doublets (Fig.8A-D) and triplets (Fig.8E-H) on a ring. In these configurations, unlike the case of linear doublets and triplets analyzed above, there are no edge cells. The critical line of *a*_*c*_ which marked the transition from CIL to CL-dominated regimes before, does not appear anymore, and there is only a very gradual shift in the behavior between the two regimes. Comparing the linear (Fig.4A,B) and ring (Fig.8B,C) pairs, we find that the ring maintains a polarized behavior at much higher values of the CIL parameter *a*, indicating that by closing the cells into a ring the polarized configuration is significantly enhanced (Fig.8D).

A similar, although more complex, behavior is observed for ring triplets (Fig.8E-H). The polarized configuration extends over a larger region of the parameter space, into larger values of the CIL parameter (compare Fig.8F and Fig.5B,G), and there is no sharp transition line between the CIL and CL regions. The stronger polarization for the ring compared with the linear train is shown in Fig.8H. Note however, that due to the inhibitory effect of the CIL interaction, the overall speed of the ring is somewhat smaller than that of the linear train, since the edge cells of the linear train do not have less CIL interactions that inhibit their motility.

Finally, when we compare long rings (Fig.8I), we find very similar behavior to the long linear trains (compare Fig.8J-L with Fig.7F-H). For small rings (*N* = 3, 4) the rings are slightly more ordered in the polarized states compared to linear trains (Fig.8M), although slower moving (Fig.8N). As the linear and ring trains grow longer than the mean size of the correlated cluster, *∼* 14 (Fig.8M), they both converge to the same dynamics (Fig.8N).

Our results therefore indicate some tendency of rings to maintain a more persistent correlated state, compared to linear trains, but this tendency diminishes with the length of the cellular cluster. The super-persistent motion of long rings of cells (*N* 40) observed in [13] are therefore not explained by our model, and may require some modification. We note that the lateral confinement of cells may influence their dynamics and cell-cell interactions across the width of the confined tracks, in a way that we do not describe in our simple one-dimensional model. In the absence of strong confinement, a finite correlation length for polarized multi-cellular motion was observed [31]. In addition, cells deposit long-lived physio-chemical footprints along their way (such as fibronectin and laminin), which determine their future migration path [15]. This effect can also contribute to the super persistence rotation observed in the experiments, which is absent from the model.

## DISCUSSION

This work presents a model for one-dimensional collective cell migration, which is based on our recently developed model for the one-dimensional motion of single cells [19]. The model contains a simple, yet explicit, mechanism for the spontaneous emergence of polarization of the individual cells in the train. This allows us to describe how the contact interactions between the cells affect their internal polarization states, which then affect the collective migration of the entire multi-cellular aggregate (train). The model predicts the configuration phase-space of cell doublets, triplets and long trains of cells, as well as rings of cells, with respect to the parameter space of the CIL and CL interactions.

The model shows how the configuration space is modified for different levels of noise, where for low values of noise the system exhibits multi-stability with many co-existing configurations of cellular polarization directions and amplitudes that arise due to the CIL and CL interactions. Within these multi-stable regions, individual cells in the trains can assume different strengths of their traction forces and internal polarization, while having the same overall polarized configuration. This result is an inherent property of the UCSP spontaneous-polarization model and provides a rich variety of multi-cellular dynamics, which goes beyond current available polarization models which treat the cells as pre-polarized SPPs with ‘Viscek-like’ interactions [13, 16].

We also demonstrate that when the noise level is sufficiently large (considering it to be more realistic), many of the transient phases lose their stability and two main configurations span the configuration space. We classify these configurations as CIL and CL by the behavior of the cells at the edges of the train, irrespective of the train size. We find that for MDCK cell doublets and triplets the proportions of polarization configurations are similar to simulations along the transition line between the CIL and CL regions. This suggests that in these cells the two types of cell-cell interactions are of similar magnitude. The model also predicts that as the multi-cellular aggregate size increases the speed and persistence decreases, as the bulk part of the train breaks down into alternating domains of opposite polarization.

In the rings configuration, our model predicts increased persistent correlated motion, similar though less dramatic compared to the experimental observations [13]. One prominent mechanism that stabilizes and drives the proliferation of domain walls in our model, is the formation of a-CIL configurations: a non-polar cell is “trapped” and depolarized by its neighbors that extend cryptic lamellipodia beneath it. These domain walls break the coherence of long cellular trains into small domains, both for linear and closed rings. Cellular mechanisms that destabilize these a-CIL configurations may explain the long-range coherence observed in experimental rings [13].

The domain walls in our model do not interact with each other very strongly, forming and diffusing independently along the cellular trains (Fig.7H,I). This is in agreement with spin-based models of one-dimensional cellular clusters, and experimental observations [16]. However, in our model each cell’s polarization can have different amplitudes, and not just a single amplitude spin, as in the spin-based models. This additional degree of freedom, which is strongly coupled between neighbors due to the CIL and CL interactions, gives rise to more complex dynamics and cellular domain configurations.

Our model demonstrates the richness of collective cell migrations, even within one-dimensional geometries, using models that explicitly account for the internal polarization state of the cells. Extending this model to flexible cells [19], will allow for more detailed comparisons with experiments.

