## Supplementary information file for "Polarization and motility of one-dimensional multi-cellular trains"

Jonathan E. Ron[1], Joseph D'alesandro[2], Victor Cellerin[2], Raphael voituriéz[3], Benoit Ladoux[2] and Nir S. Gov[1]
[1] *Department of Chemical and Biological Physics, Weizmann Institute of Science, Israel*
[2] *Université de Paris, CNRS, Institut Jacques Monod, F-75006 Paris, France and*
[3] *Laboratoire Jean Perrin and Laboratoire de Physique Thorique de la Matire Condense, Sorbonne Universit, Tour 13-12, 5eme etage, 4 place Jussieu, 75252 Paris Cedex 05, France*

### APPENDIX A - MODEL DERIVATION FOR A SINGLE CELL

Consider a cell with a fixed length of $l$. The edges of the cell are denoted by $x_f$ and $x_b$ (front an back), and at each edge actin polymerizes, and produces local actin flows with a mean speed of $v_f$ and $v_b$ respectively. The global actin flow inside the cell, or the net retrograde flow, is given by the difference between the actin polymerization speeds at the two edges

$$v = v_f - v_b \tag{S.1}$$

Note that in this model the local polymerization speeds (at which actin polymerizes towards the edge) and the local actin flows (which flow retrograde inward the center) are considered to be of the same magnitude, and from hence forward we will consider the local actin activity $v_f$ and $v_b$ via the polymerization speeds.

Consider now a chemical cue, which inhibits actin polymerization at the edges of the cell near the membrane. Such inhibitory cues can be associated with capping proteins, which cap the edges of the polymerizing actin filaments and prevent their elongation [1], or proteins such as arpin which inhibit formation of arp2/3 nucleation sites near the membrane [2, 3]. We note the chemical cue by $c(x)$.

The chemical cue inside the cell follows an advection-diffusion transport: Along a segment of $\delta x$ it diffuses with a typical time of $\tau_D = \delta x^2 D^{-1}$ ($D$ is the diffusion coefficient), and is advected with the actin flow at the velocity of the net flow $v$. The typical time of the actin flow to pass a segment of $\delta x$ is therefore $\tau_v = \delta x v^{-1}$.

We denote the the typical time scale at which the polymerization speed $v_i$ changes by $\tau_p$, and assume that $\tau_p \gg \tau_D, \tau_v$, i.e, the chemical cue has already diffused and advected and the molecules are ready to react with the actin actin filaments before any changes in the polymerization speeds have occurred [3, 4].

We consider the polarity cue chemical which is advected with the net retrograde flow of actin, and only at the cell's edge it associates and affects the actin polymerization. We write the advection-diffusion transport $c(x)$ at a steady state, i.e, after the distribution of the chemical cue along the cell (the segment $l$) reached equilibrium (as done in [3, 4])

$$0 = \frac{dx}{dt}\left(vc(x) + D\frac{dc(x)}{dt}\right) \tag{S.2}$$

The solution of (S.2) reads

$$c(x) = c_0 exp\left(-\frac{vx}{D}\right) + c_1 \tag{S.3}$$

and the coefficients are obtained by applying a no-flux boundary condition at $x_f$ and $x_b$

$$0 = vc(x) + D\left.\frac{dc(x)}{dx}\right|_{x=x_b} \tag{S.4}$$

$$0 = vc(x) + D\left.\frac{dc(x)}{dx}\right|_{x=x_f} \tag{S.5}$$

We next assume the the total amount of the chemical cue is conserved inside the cell throughout the process

$$c_{tot} = c_0\int_{x_b}^{x_f} e^{-\frac{vx}{D}}dx \rightarrow c_0 = \frac{c_{tot}v}{D}\left(\frac{1}{e^{-\frac{vx_b}{D}} - e^{-\frac{vx_f}{D}}}\right) \tag{S.6}$$

The full exponential form of the concentration profile along the cell is given by inserting (S.6) into (S.3)

$$c(x)=\frac{c_{tot}v}{D}\left(\frac{exp\left(-\frac{vx}{D}\right)}{1-exp\left(-\frac{vl}{D}\right)}\right) \tag{S.7}$$

We will now address reaction kinetics of the chemical cue with the active actin filaments at the cell's edges, noted by $n_f$ and $n_b$.

We write the binding and unbinding dynamics of $n_i$ in the presence of the inhibitory cue $c(x_i)$ ($i$ notes the front or the back), which acts as a capping protein of the polymerizing actin filaments

$$\frac{dn_i}{dt}=k^c_{off}(1-n_i)-k^c_{on}n_ic(x_i) \tag{S.8}$$

where $k^c_{on}$ and $k^c_{off}$ are the kinetic rates of the inhibitory ("capping") reaction.

At steady state, the proportion of the actin filaments is given by

$$n_i=\frac{c_s}{c_s+c(x_i)} \tag{S.9}$$

where $c_s=\frac{k^c_{off}}{k^c_{on}}$.

Utilizing the UCSP model, we write the polymerization speeds at both ends of the cell as

$$v_i=\beta\left(\frac{c_s}{c_s+c(x_i)}\right) \tag{S.10}$$

and therefore the net retrograde flow of actin for a cell of length $l$ at steady state where the boundaries are $x_b=0$ and $x_f=l$ is given by

$$v=\beta\left(\frac{c_s}{c_s+c(0)}-\frac{c_s}{c_s+c(l)}\right) \tag{S.11}$$

which we write for in a compact way throughout the paper as

$$v^*=\beta\left(\frac{1}{1+c_0}-\frac{1}{1+c_L}\right) \tag{S.12}$$

where $c_0, c_L$=$\frac{c(0)}{c_s}$, $\frac{c(l)}{c_s}$

We derive now the time evolution equation for the polymerization speed, noting that the kinetic rates $k_{on}$ and $k_{off}$ define the time scale $\tau_p$, i.e, as changes in the inhibitory cue concentration affect the proportion of the active polymerized actin and therefore the speed of polymerization instantaneously (Eq. S.8).

Considering that $v_i=\beta n_i$ (S.10) we can write (S.8) as

$$\begin{aligned}\frac{dv_i}{dt} &= k^c_{off}(\beta-v_i)-v_ik^c_{on}c(x_i) && \text{(S.13)}\\ &= \beta k^c_{off}-v_i\left(k^c_{off}+k^c_{on}c(x_i)\right) && \text{(S.14)}\end{aligned}$$

we add a small perturbation to $v$ from its steady state: $v\approx v^*_i+\epsilon$ and obtain

$$\begin{aligned}\frac{d\epsilon}{dt} &= \beta k^c_{off}-v^*_i\left(k^c_{off}+k^c_{on}c(x_i)\right)-\epsilon\left(k^c_{off}+k^c_{on}c(x_i)\right) && \text{(S.15)}\\ &= -\epsilon\left(k^c_{off}+k^c_{on}c(x_i)\right) && \text{(S.16)}\end{aligned}$$

substituting back $\epsilon\approx v_i-v^*_i$ we obtain

$$\frac{dv_i}{dt}=-\delta\left(v_i-v^*_i\right) \tag{S.17}$$

where $\delta=\left(k^c_{off}+k^c_{on}c(x_i)\right)=k^c_{on}(c_s+c(x_i))$. For simplicity, we treat $\delta$ as a constant in our analysis, neglecting the time-dependence due to dynamic changes in $c(x_i)$.

### APPENDIX B - MODEL CALIBRATION FOR SINGLE CELL PARAMETERS

We consider the rest length of the cell to be $l_0 = 30$ [µm], and that the diffusion coefficient is in the order of magnitude of $D \sim$ 1-10 $\left[\frac{\mu m^2}{s}\right]$.

Using these estimation we can evaluate the adhesion turnover rate

$$k^0_{off} = \frac{D}{dl_0^2} = \frac{1-10}{4\times 30^2} = 2.8\times 10^{-4} - 2.8\times 10^{-3} \approx 10^{-4} - 10^{-3}\left[s^{-1}\right] \tag{S.18}$$

and therefore the coupling parameter can be estimated by

$$\beta = \frac{v}{l_0 k^0_{off}} = \frac{0.1}{30\times(10^{-4}-10^{-3}} = 3.33 - 33.3 \approx 3 - 30 \tag{S.19}$$

The data shows that $v_{cell} \approx 50\left[\frac{\mu m}{hr}\right] = 0.015\left[\frac{\mu m}{s}\right]$, therefore the normalized velocity is evaluated by

$$\tilde{v}_{cell} = \frac{v_{cell}}{c_0 k^0_{off}} = \frac{0.015}{30\times(10^{-4}-10^{-3})} \approx 0.5 - 5 \tag{S.20}$$

The data also shows that for a persistent cell the average length is $l = 62.25$ [µm], therefore

$$\tilde{l} = \frac{l}{l_0} = \frac{62.5}{30} \approx 2.1 \tag{S.21}$$

The fitting procedure is shown in Fig.S.1.

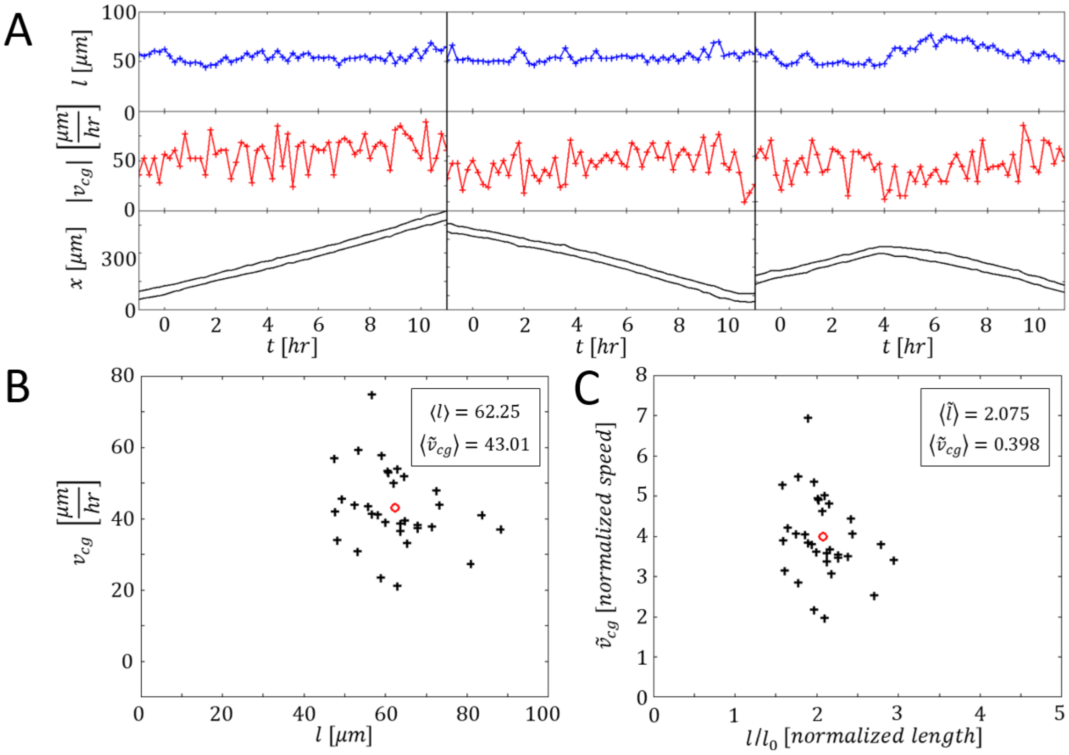


FIG. S.1: A) Kymographs of a single MDCK cells migrating smoothly with a fixed length. Upper/Middle/Lower panel indicate the time series of the cell length/cell speed/kymograph. B) Scatter plot of the average cell length and speed. C) Scatter plot of the normalized average cell length and speed. The speed is normalized by $\tilde{v}_{cg} = \frac{v_{cg}}{k^0_{off} l_0}$, where $l_0 = 30$ [µm] (rest length), and $k^0_{off} = 10^{-4}\left[s^{-1}\right]$ (adhesion turnover rate). $n = 32$. Red circles indicate the average.

### APPENDIX C - FOKKER PLANK SOLUTION FOR CELL PAIRS

The stochastic differential equations of the cell pair can be written as a Langevin equations

$$dv_i = \delta F_i(v_i)dt + \sigma dW_i \tag{S.22}$$

where $\delta$ is the damping rate and $dW_i = \xi_t\sqrt{dt}$ (Wiener process), and $\xi_t$ is a Gaussian random noise with an amplitude of $\sigma$. The index $i$ denotes the number of the cell, i.e, $i = 1, 2$.

The steady-state distributions of $v_1$ and $v_2$ can be given by the solution of the Fokker-Plank equations derived for (Eqs.11-13 in main text)

$$\frac{\partial p_i\left(v_i, t\right)}{\partial t} = -\delta\frac{\partial\left(F_i(v_i)o_i(v_i,t)\right)}{\partial v_i} + \frac{\sigma^2}{2}\frac{\partial^2 p_i(v_i)}{\partial v_i^2} \tag{S.23}$$

where at steady-state $p(v_i, t \to \infty) = p(v_i)$ and therefore

$$\frac{\partial\left(F_i(v_i)p_i(v_i)\right)}{\partial v_i} = \frac{\sigma^2}{2\delta}\frac{\partial^2 p_i(v_i)}{\partial v_i^2} \to \frac{\partial}{\partial v_i}\left(\underbrace{F_i(v_i)p_i(v_i) - \frac{\sigma^2}{2\delta}\frac{\partial p_i(v_i)}{\partial v_i}}_{\equiv C_0}\right) = 0 \tag{S.24}$$

Since the distribution is normalizeable $p_i\left(v_i \to \pm\infty\right)$ then $c_0 = 0$, and the solution for Eq.(S.24) is given by

$$p_i(v) = \frac{exp\left(\frac{2\delta}{\sigma^2}\int_0^{v_i} F_i(u)du\right)}{\int_{-\infty}^{+\infty} exp\left(\frac{2\delta}{\sigma^2}\int_0^{v} F_i(u)du\right)dv} \equiv C_i exp\left(-W_i\left(v_i\right)\right) \tag{S.25}$$

where the constant of integration at the denominator is given by the normalization condition $\int_{-\infty}^{+\infty p_i(v_i)dv_i} = 1$ and $W(v)$ is the effective energy potential. The distribution of the center-of-mass velocity $v_{cm} = \frac{v_1+v_2}{2}$ is calculated by convolving $p_1(v)$ and $p_2(v)$ in the following manner

$$p_{cm}(v) = 2\int_{-\infty}^{+\infty} p_1(2v - w)p_2(w) \tag{S.26}$$

where we have used the following Fourier transform identities

$$p(v) = \int_{-\infty}^{+\infty} p_1(v-u)p_2(u)du \tag{S.27}$$

$$F\left[p(v)\right] = P_1(k)P_2(k) \tag{S.28}$$

$$F\left[p\left(\frac{v}{2}\right)\right] = 2H(2k) \tag{S.29}$$

$$p\left(\frac{v}{2}\right) = 2\int_{-\infty}^{+\infty} p_1(2v-u)p_2(u)du \tag{S.30}$$

We find that the calculations are accurate for $q = 0$ and $q \gg 1$, however for small finite values of $q$, the joint distribution is required. The results are shown in Fig.S.2

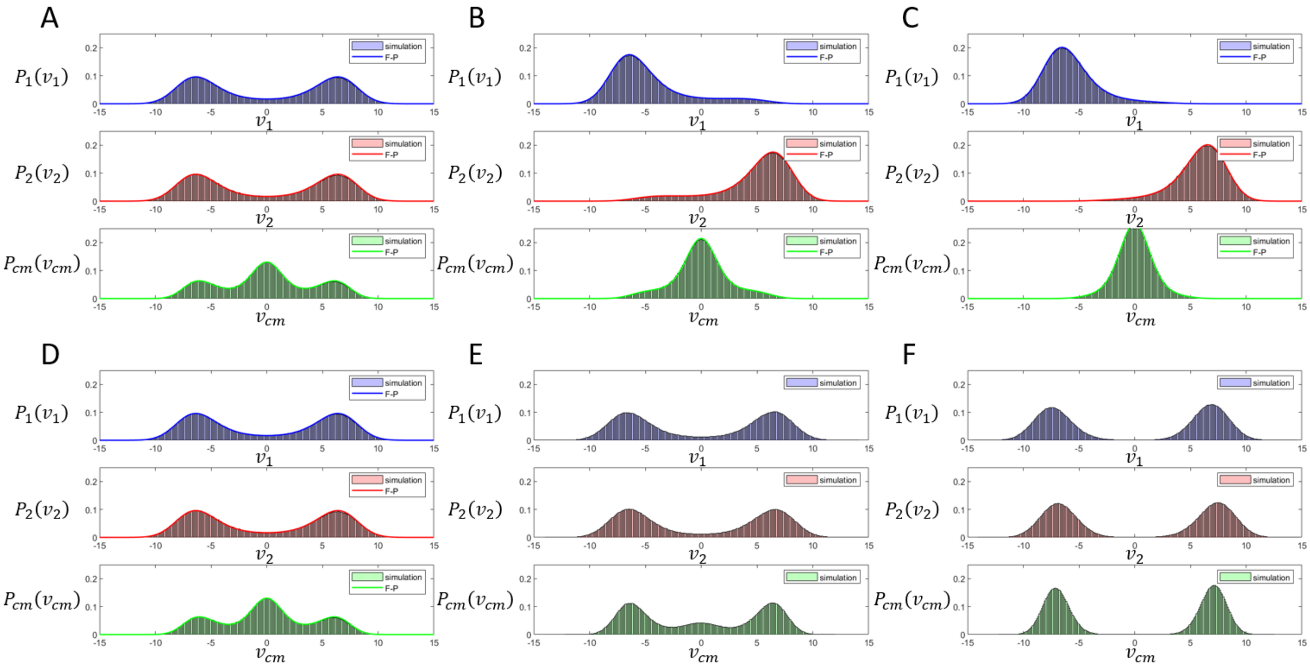


FIG. S.2: Simulations and their corresponding Fokker-Plank solutions. A-C) The steady-state distribution dynamics with an increase of the CIL interaction .D-F) The steady-state distribution dynamics with an increase of the CL interaction. Blue/Red/Green stand for $v_1/v_2/v_{cm}$ respectively.

### APPENDIX D - STEADY STATE ANALYSIS - CELL PAIRS

To depict the $a-q$ phase diagram we analyze the steady state solutions of $v_1$ and $v_2$ (Eqs. 9-11 in the main text) by solving the following functionals

$$F_1(v) = v - \beta\left(\frac{1}{1+a+q\cdot c_L(v)} - \frac{1}{1+c_0(v)}\right) \rightarrow F_1(v) = 0 \tag{S.31}$$

$$F_2(v) = v - \beta\left(\frac{1}{1+c_L(v)} - \frac{1}{1+a+q^{-1}c_0(v)}\right) \rightarrow F_2(v) = 0 \tag{S.32}$$

where here for the purpose of the analysis the CL effect is constrained into a single direction.

The functionals (Eqs.S.31,S.32) show that when no interactions are introduced both cells act as independent UCSP cells which can polarize in either direction freely irrespective to one another (Fig.S.3A). When a CIL interaction is introduced ($q=1$ and $a>0$), both functionals have a single stable solution which is symmetric and opposite in sign, corresponding to two cells which polarize away from each other (Fig.S.3B). When a CL interaction is introduced ($q>1$ and $a=0$) both functionals have a single stable solution of the same sign but with a different magnitude, which corresponds to a polarized pair in which the cell in the front has a faster global internal actin flow (Fig.S.3C).

In this analysis we also find the $a_c$ transition line (Black solid line in Fig.3A in the main text) by taking the limit of $v \rightarrow 0$ in $F_2(v)$ (the winning cell)

$$\lim_{v\rightarrow 0}\{F_2(v)\} = \beta\left(\frac{1}{1+\frac{c}{l}} - \frac{1}{1+a+q^{-1}\frac{c}{l}}\right) \rightarrow a_c = \frac{c}{l}\left(1-q^{-1}\right) \tag{S.33}$$

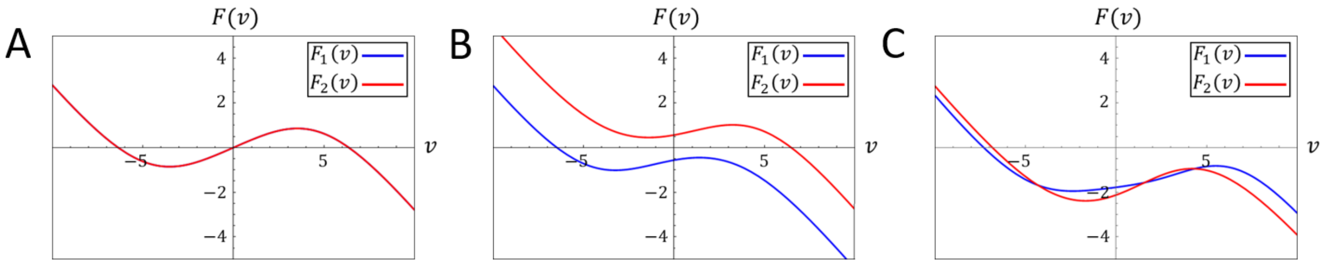


FIG. S.3: The functional forms (Eqs.S.31,S.32 for a configuration in which cell 2 is the winning cell). Blue/Red curves indicate $F_1(v)/F_2(v)$ respectively. Solutions for: A) $a=0$ and $q=1$. B) $a>0$ and $q=1$. C) $a=0$ and $q>1$.

### APPENDIX E - EXTENDED STEADY-STATE ANALYSIS OF THE DOUBLET SYSTEM

In this section we present an extensive analysis of the doublet dynamics along the different sections of the $a$-$q$ phase-diagram (Fig.3 in main text) for low and large values of noise.

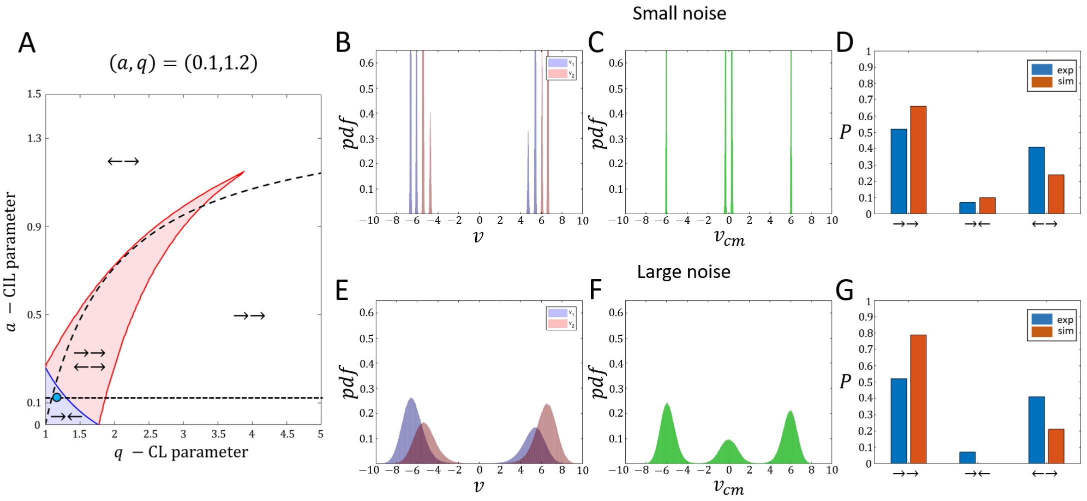


FIG. S.4: Steady-state behavior of a doublet system with $(a, q) = (0.1, 1.2)$. A) The $a$-$q$ phase-diagram indicating the analyzed point (teal circle) along the $a$ cross-section (dashed black horizontal line). B-D) Steady-state dynamics for a small level of noise ($\sigma = 1/32$). E-F) Steady-state dynamics for a large level of noise ($\sigma = 1/\sqrt{2}$). Blue/Red/Green distributions indicate the $v_1/v_2/v_{cm}$ probability density functions respectively. Blue/Orange bars indicate the frequency of appearance for the configurations as measured in the experiments/simulations respectively.

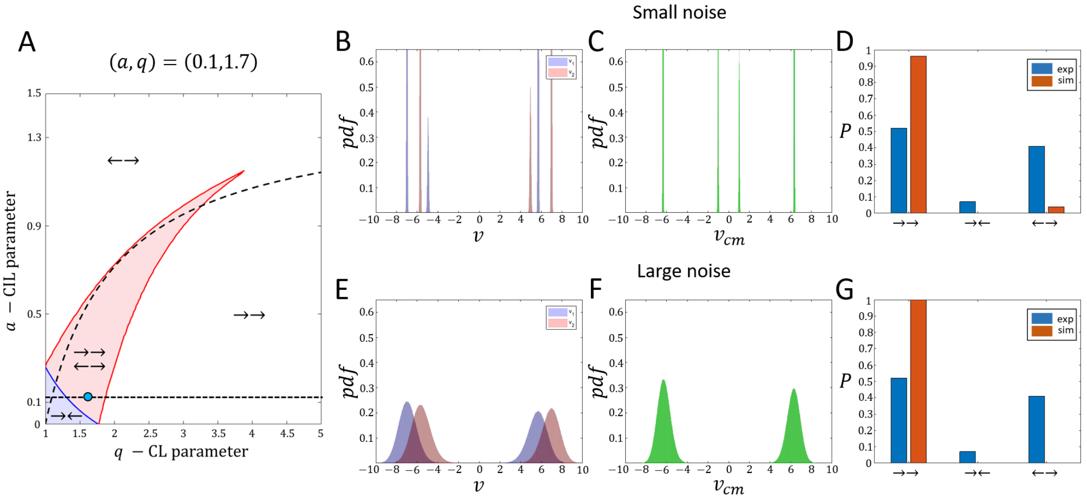


FIG. S.5: Steady-state behavior of a doublet system with $(a, q) = (0.1, 1.7)$. A) The $a$-$q$ phase-diagram indicating the analyzed point (teal circle) along the $a$ cross-section (dashed black horizontal line). B-D) Steady-state dynamics for a small level of noise ($\sigma = 1/32$). E-F) Steady state dynamics for a large level of noise ($\sigma = 1/\sqrt{2}$). Blue/Red/Green distributions indicate the $v_1/v_2/v_{cm}$ probability density functions respectively. Blue/Orange bars indicate the frequency of appearance for the configurations as measured in the experiments/simulations respectively.

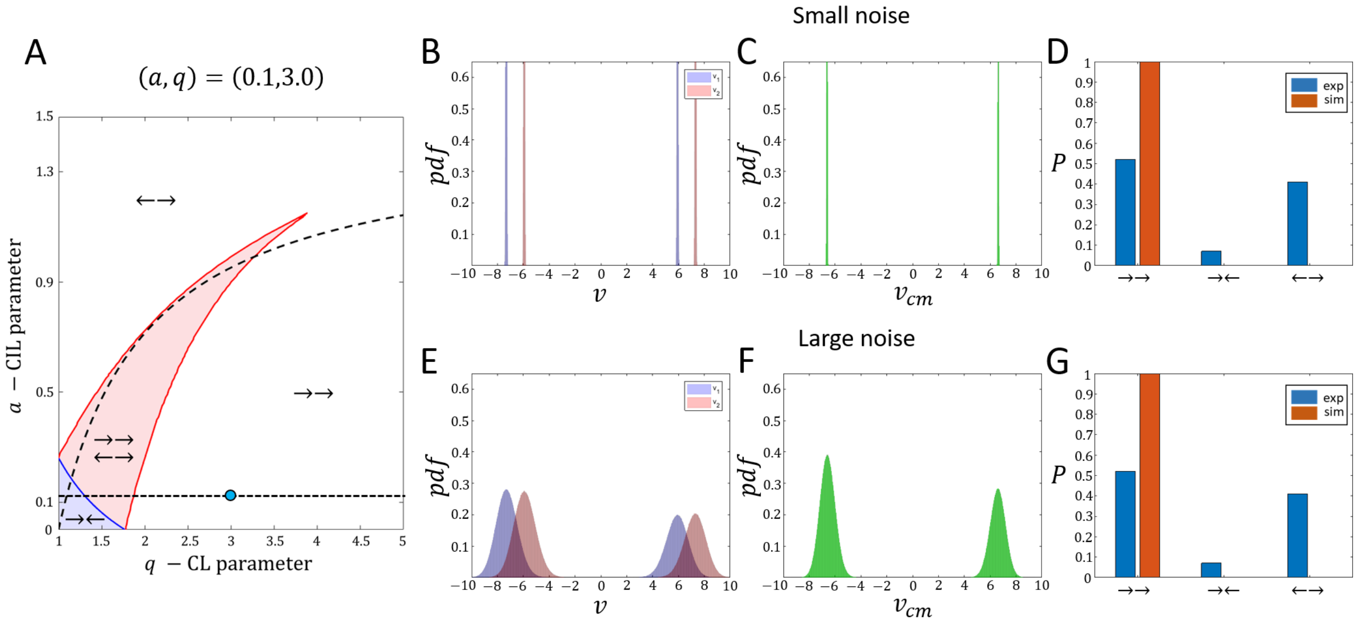


FIG. S.6: Steady-state behavior of a doublet system with $(a, q) = (0.1, 3)$. A) The $a$-$q$ phase-diagram indicating the analyzed point (teal circle) along the $a$ cross-section (dashed black horizontal line). B-D) Steady-state dynamics for a small level of noise ($\sigma = 1/32$). E-F) Steady-state dynamics for a large level of noise ($\sigma = 1/\sqrt{2}$). Blue/Red/Green distributions indicate the $v_1/v_2/v_{cm}$ probability density functions respectively. Blue/Orange bars indicate the frequency of appearance for the configurations as measured in the experiments/simulations respectively.

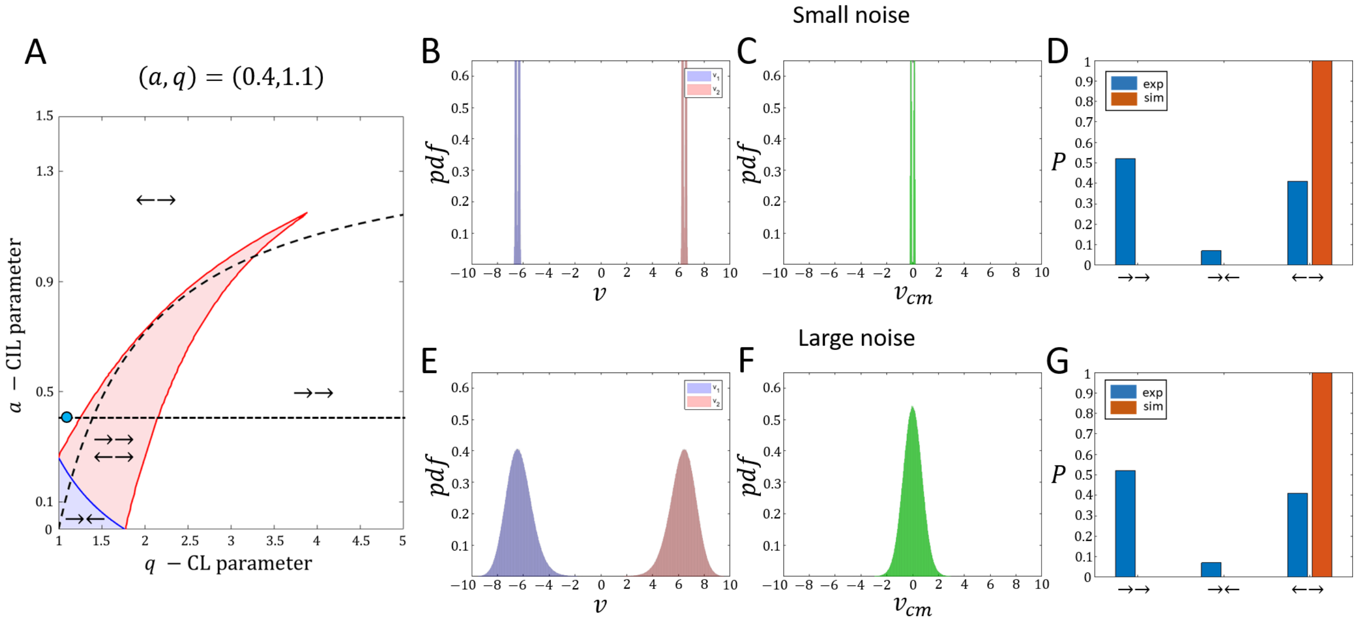


FIG. S.7: Steady-state behavior of a doublet system with $(a, q) = (0.4, 1.1)$. A) The $a$-$q$ phase-diagram indicating the analyzed point (teal circle) along the $a$ cross-section (dashed black horizontal line). B-D) Steady-state dynamics for a small level of noise ($\sigma = 1/32$). E-F) Steady-state dynamics for a large level of noise ($\sigma = 1/\sqrt{2}$). Blue/Red/Green distributions indicate the $v_1/v_2/v_{cm}$ probability density functions respectively. Blue/Orange bars indicate the frequency of appearance for the configurations as measured in the experiments/simulations respectively.

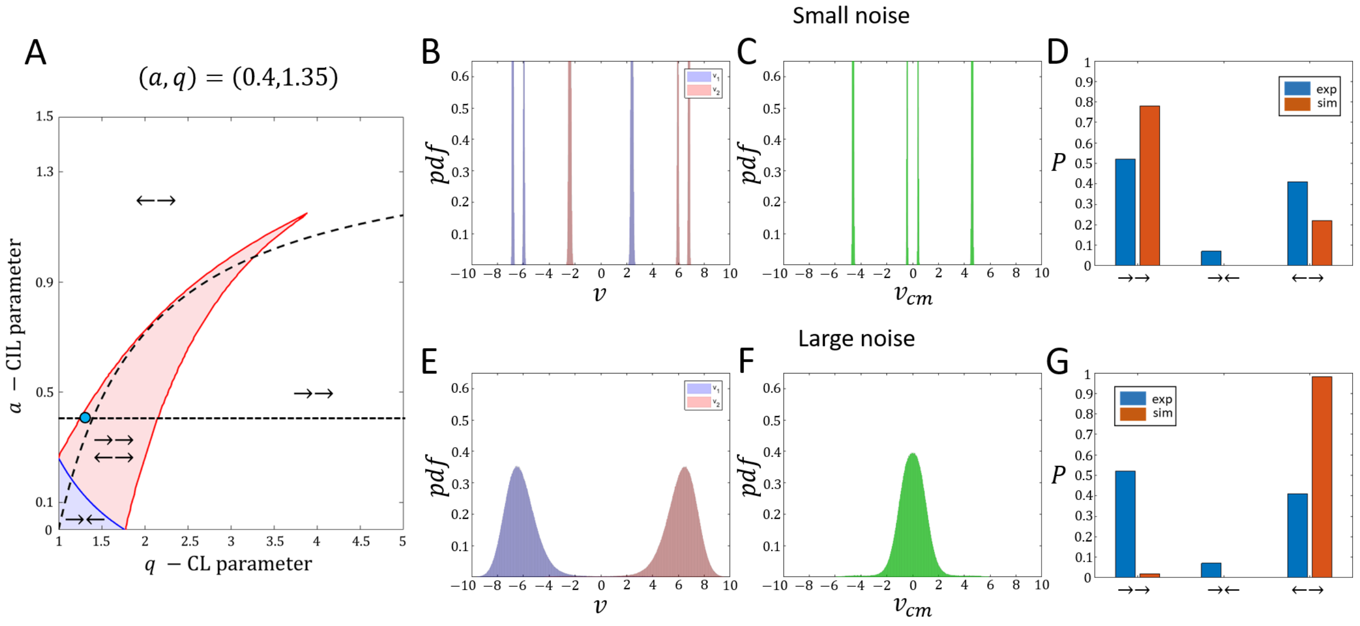


FIG. S.8: Steady-state behavior of a doublet system with $(a, q) = (0.4, 1.35)$. A) The $a$-$q$ phase-diagram indicating the analyzed point (teal circle) along the $a$ cross-section (dashed black horizontal line). B-D) Steady-state dynamics for a small level of noise ($\sigma = 1/32$). E-F) Steady-state dynamics for a large level of noise ($\sigma = 1/\sqrt{2}$). Blue/Red/Green distributions indicate the $v_1/v_2/v_{cm}$ probability density functions respectively. Blue/Orange bars indicate the frequency of appearance for the configurations as measured in the experiments/simulations respectively.

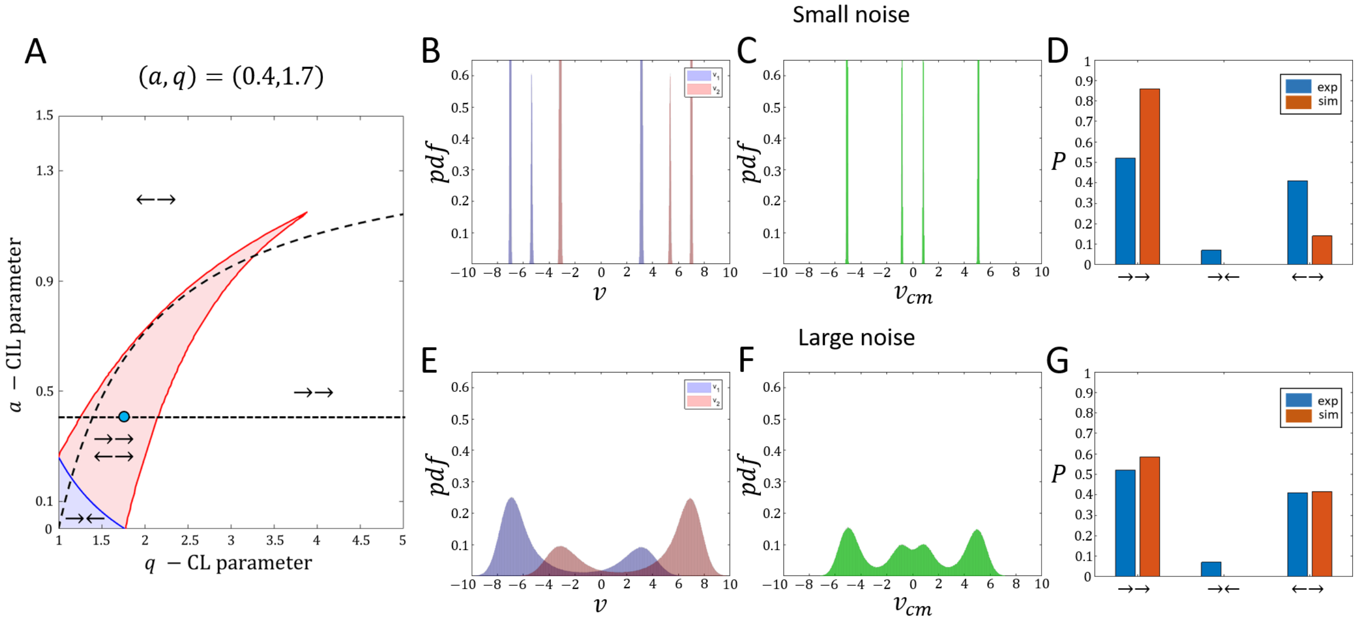


FIG. S.9: Steady-state behavior of a doublet system with $(a, q) = (0.4, 1.7)$. A) The $a$-$q$ phase-diagram indicating the analyzed point (teal circle) along the $a$ cross-section (dashed black horizontal line). B-D) Steady-state dynamics for a small level of noise ($\sigma = 1/32$). E-F) Steady-state dynamics for a large level of noise ($\sigma = 1/\sqrt{2}$). Blue/Red/Green distributions indicate the $v_1/v_2/v_{cm}$ probability density functions respectively. Blue/Orange bars indicate the frequency of appearance for the configurations as measured in the experiments/simulations respectively.

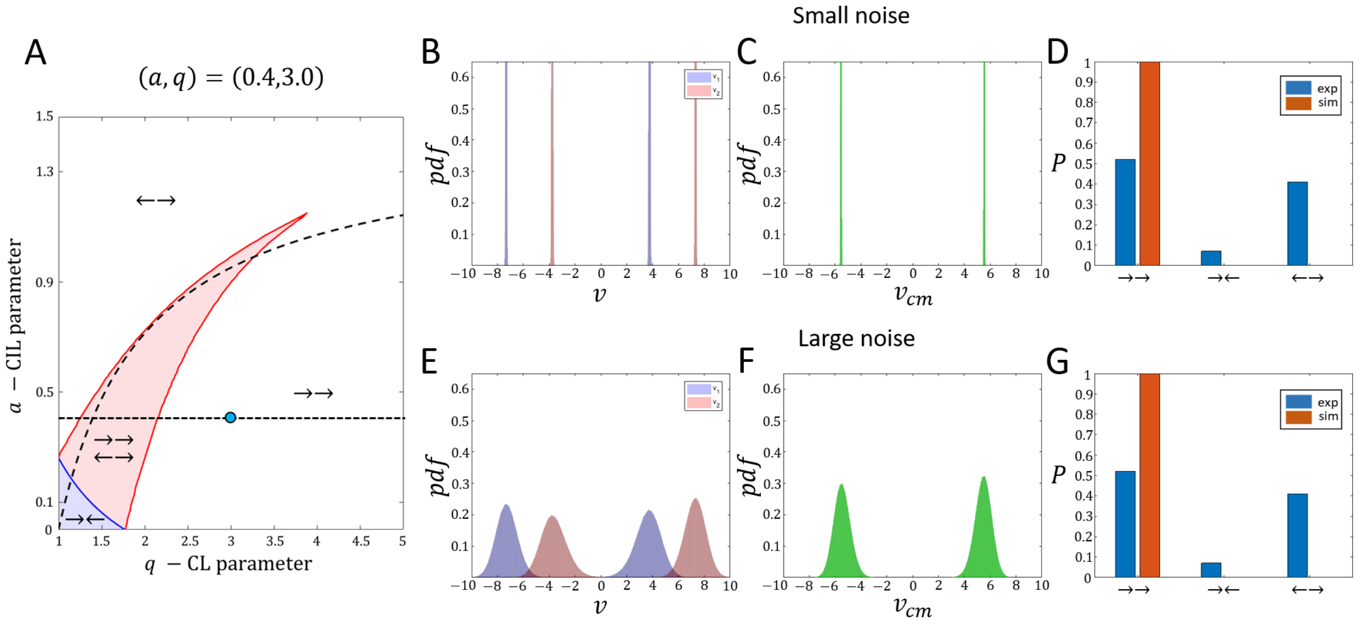


FIG. S.10: Steady-state behavior of a doublet system with $(a, q) = (0.4, 3)$. A) The $a$-$q$ phase-diagram indicating the analyzed point (teal circle) along the $a$ cross-section (dashed black horizontal line). B-D) Steady-state dynamics for a small level of noise ($\sigma = 1/32$). E-F) Steady-state dynamics for a large level of noise ($\sigma = 1/\sqrt{2}$). Blue/Red/Green distributions indicate the $v_1/v_2/v_{cm}$ probability density functions respectively. Blue/Orange bars indicate the frequency of appearance for the configurations as measured in the experiments/simulations respectively.

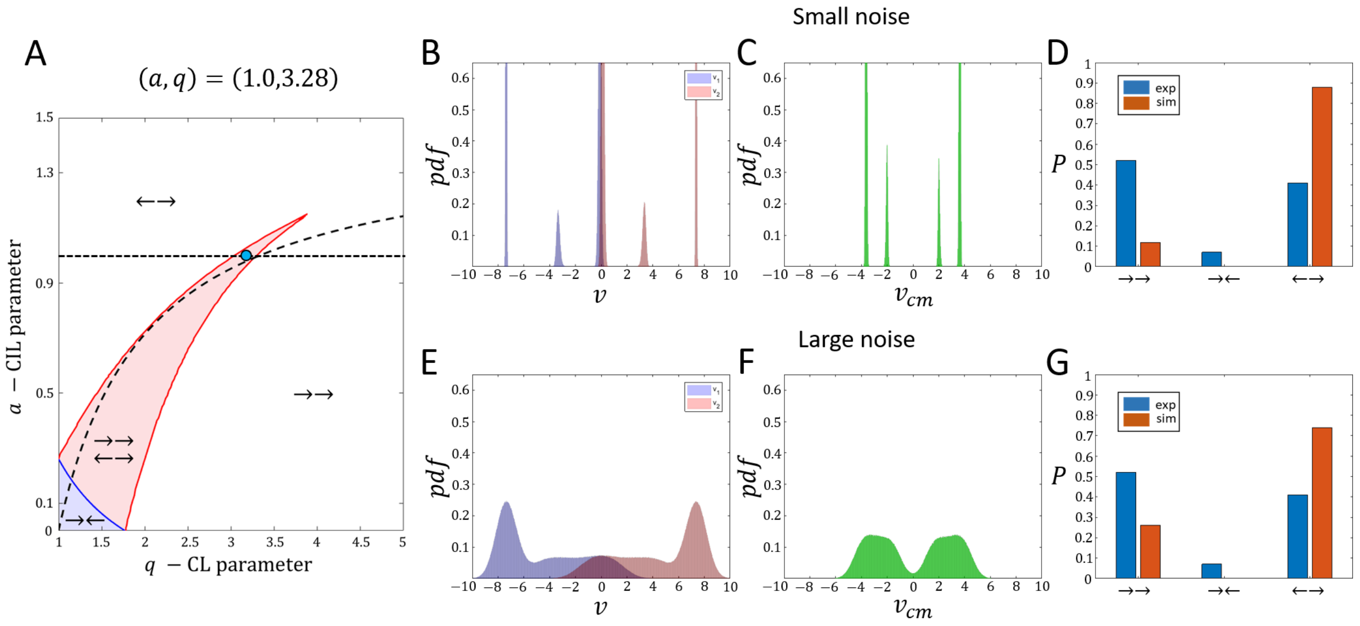


FIG. S.11: Steady-state behavior of a doublet system with $(a, q) = (1, 3.28)$. A) The $a$-$q$ phase-diagram indicating the analyzed point (teal circle) along the $a$ cross-section (dashed black horizontal line). B-D) Steady-state dynamics for a small level of noise ($\sigma = 1/32$). E-F) Steady-state dynamics for a large level of noise ($\sigma = 1/\sqrt{2}$). Blue/Red/Green distributions indicate the $v_1/v_2/v_{cm}$ probability density functions respectively. Blue/Orange bars indicate the frequency of appearance for the configurations as measured in the experiments/simulations respectively.

### APPENDIX F - EXPERIMENTAL METHODS

**Cell culture**
We used MDCK WT, MDCK histon-GFP and MDCK PBD-YFP (gift from F. Martin-Belmonte lab). The cells were cultured in DMEM GlutaMAX high-glucose (Gibco, Waltham, MA) supplemented with 10 % foetal bovine serum (BioWest, Nuaillé, France). Prior to experiments, the cells were treated with mitomycin C at final concentration of 10 µg/mL added in the medium for 1 h, then rinsed before subsequent detachment and seeding on the experimental samples.

**Sample preparation**
All micropatterns were prepared using standard micro-contact printing on PDMS, as described in Ref. [5]. The substrates used were: (i) non-culture treated plastic dishes (Greiner Bio-One, Kremsmünster, Austria) for wide-field microscopy or (ii) glass coverslips (Menzel-Gläser) for spinning-disk microscopy. The substrates were first covered with a thin layer of poly-dimethylsyloxane (PDMS, Sylgard, Dow Corning, Midland, MI) using a spin-coater and crosslinked at 80°C for 2 h. PDMS stamps were made by pouring PDMS on a mold featuring the patterns to be printed and crosslinked as described. After cooling down, a fibronectin solution was prepared by adding 50 $\mu$g/mL of fibronectin and 25 $\mu$g/mL of Cy3- or Cy5-labelled fibronectin into sterile milliQ water. The solution was then incubated on the stamps for 40 min at room temperature. Before stamping, the substrates were activated using UV-ozone for 10 min; the stamps were rinsed to remove any excess fibronectin and dried using an air-gun. The stamps were briefly put in contact with the surface of the substrate, then removed, and the substrates immerged in a 2% pluronics F127 (Sigma-Aldrich, Saint Louis, MO) solution in PBS for 2 h. Finally, the substrates were rinsed in PBS and sterilized under the UV lamp of a culture hood before use.

**Cell seeding**
The cells were enzymatically detached, then concentrated using a centrifuge and seeded on the substrates in culture medium, at a controlled density: medium-low density for the doublet and random small trains experiments, very high density for the stencil experiments, a wide range of densities for the ring experiments. The cells were let to adhere in the incubator for approx. 45 min, then rinsed thoroughly (but carefully) to remove excess floating cells without affecting adhered cells.

**Time-lapse microscopy**
All experiments were run at 37°C in 5% CO2. The experiments with PBD-YFP cells were done using an inverted microscope (Leica, Wetzlar, Germany) with a CSU-W1 confocal spinning-disk module (Yokogawa, Tokyo, Japan) and a 40X oil-immersion objective. The acquisition was done using Metamorph (Molecular Devices, San Jose, CA), at a 6 to 10 minutes acquisition-rate. The focus was done on the basal plane of the cells and the microscope's hardware autofocus was used to ensure the absence of defocusing. All the other experiments were done with a wide-field inverted microscope (Olympus, Tokyo, Japan) using a 10X air objective. Phase contrast and GFP fluorescence images were acquired using Metamorph (Molecular Devices, San Jose, CA), at a 6 to 12 minutes acquisition-rate. An image of the labelled patterns was done at least at the beginning of the experiment to allow further alignment.

**Polarity measurements**
For polarity measurements, the images were first visually inspected using Fiji to locate cell doublets, then the images were rotated, cropped and stitched using in-house programs in Fiji and Matlab (The Mathworks, Natick, MA). The polarities were retrieved both using a semiautomated and a manual procedure due to the intrinsic heterogeneity in PBD-YFP signal. The semiautomated measurements involved drawing a rectangular ROI at each frame for each cell in order to deal with (i) spurious, high intensity signals found mainly at the cell edges and (ii) the difficulty of precisely detecting cell edges. The PBD-YFP intensity was then averaged in the direction transverse to the line and normalized so as to range between -0.5 and 0.5. Finally, the x-coordinate along the cell was interpolated on a normalized symmetric coordinate x and the polarity was computed as the first moment of the normalized PBD-YFP signal along those symmetric coordinates. The PBD-YFP signal heterogeneity greatly limited the efficacy of the semi-automated protocol described above and led to the correct analysis of a small number of cells. Thus, for the quantitative measurements of configurations statistics, we manually assessed the polarity of single cells, based on the visual inspection of both PBD-YFP and transmission images. The combination of the information contained in those images (PBD-YFP peaks on the edges, cell shape, lamellipodium visible in transmission) allowed us to non-ambiguously determine the – binary – cell polarities in most of the frames, while ambiguous data points were simply discarded.

**Kymograph analysis**
To generate khymographs of the PBD-YFP signal along cell trains, a rectangular selection was first made in ImageJ, spanning all the train trajectory along the line direction, and approx. 16 pixels around the midline of the line pattern

in the perpendicular direction, then the Image>Stacks>Reslice function was used to generate a stack of khymograph (akin to switching the perpendicular / vertical dimension with the time / stack depth), and the median value was taken using the Image>Stacks>Z Project function. The aspect ratio of the final khymograph was then adjusted for visualisation using the Image>Adjust>Size function with free aspect ratio and no interpolation options. The zoomed-in khymographs were generated by selecting a rectangle ROI larger than the line width and simply concatenating subsequent time frames along the vertical axis.

### APPENDIX G - PROCEDURE OF PROPORTIONS CALCULATIONS

Here we list the statistical protocol used to compute the proportions of the configurations, and the proportions of the internal states within each configuration.

**For the train configuration:**

1. choose a set of $(a, q)$ for the simulation.
2. Run $N_r = 300$ realizations of length $T = 10^5$ with a time step of $dt = 0.1$ (i.e, $N_{steps} = 10^6$).
3. For each realization calculate the local flows ($v_1, v_2, v_3$ and $v_{cm}$ for the center-of-mass velocity (Fig.S.12A).
4. Smooth the trajectories (Fig.S.12B).
5. Binarize the smoothed trajectories, such that that negative/positive velocities assume a values of $+1/-1$, and absolute values below the noise threshold assume a value of 0 (Fig.S.12C).
6. Count in the entire run the data for each configuration $N_c$, and calculate the proportion by $\frac{N_c}{N_r \times N_{Steps}}$

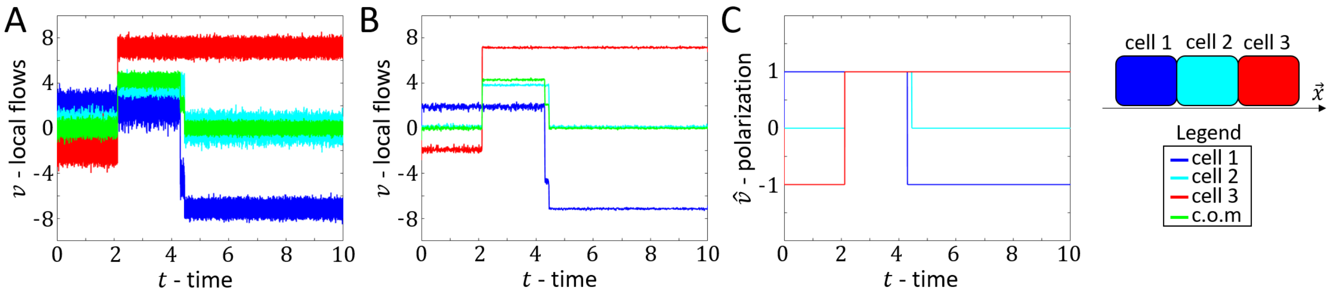


FIG. S.12: Example of the configuration classification procedure. Parameters correspond to Fig.5E in the main text. A) local flow and center-of-mass velocity trajectories. B) Smoothed trajectories. C) Binarized trajectories. Blue/Teal/Red/Green corresponds to $v_1/v_2/v_3/v_{cm}$.

**For the internal states within a configuration:**

1. Collect the center-of-mass velocities for each configuration separately.
2. For each individual configuration calculate the probability density function of the center-of-mass velocity (blue curves in Fig.S.13).
3. Within the probability density function of the center-of-mass velocity, find and label the peaks (Red circles in Fig.S.13).
4. If the number of peaks is larger than 1 in the probability density function of the center-of-mass velocity (taken for the absolute value of $v_{cm}$), insert a mid-line between two neighbouring peaks (Black dashed lines in Fig.S.13).
5. The fraction of each internal state within a configuration is calculated by the density within two neighbouring peaks, divided by the entire density function. Note that we assume here that each peak corresponds to a different internal state.

### APPENDIX H - TRIPLET SYSTEM FULL ANALYSIS

To analyze the triplet system, we investigate all the possible functional forms that the steady-state global flow in the middle cell ($v_2$ in Eq.14 in the main text) obeys

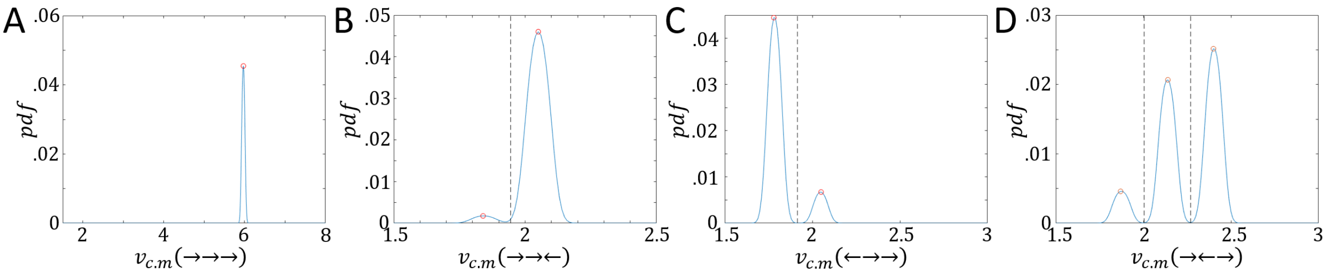


FIG. S.13: Example of the internal state classification within each configuration. Parameters correspond to Fig.5A in the main text. The probability density function for the center-of-mass velocity of configuration: A) →→. B) →→←. C) ←→→. D) →←→. Red circles correspond to the peaks. Black dashed lines correspond to the mid-line between the peaks.

$$F_{<>}(v) = -v + \beta\left(\frac{1}{1+a+q^{-1}c_L(v)} - \frac{1}{1+a+q^{-1}c_0(v)}\right) \rightarrow F_{CIL}(v) = 0 \tag{S.34}$$

$$F_{><}(v) = -v + \beta\left(\frac{1}{1+a+qc_L(v)} - \frac{1}{1+a+qc_0(v)}\right) \rightarrow F_{C\bar{I}L}(v) = 0 \tag{S.35}$$

$$F_{>>}(v) = -v + \beta\left(\frac{1}{1+a+qc_L(v)} - \frac{1}{1+a+q^{-1}c_0(v)}\right) \rightarrow F_{CL}(v) = 0 \tag{S.36}$$

where the labels of the functionals refer to the internal polarization of the middle cell (Fig.S.14A).

A qualitative study of the functionals shows that for a pure CIL interaction ($a > 0$ and $q = 1$) the middle cell can assume only a non-polarized state of $v_2 = 0$, while for a pure CL interaction ($a = 0$ and $q > 1$) several meta-stable states can co-exist (Fig.S.14B). We note that the functional analysis of the edge cells (cells 1 and 3) consists of the doublets' functionals (Eqs.S.31,S.32).

Using this analysis we construct the extended $a - q$ phase diagram for the triplet train, which is composed of all the possible configurations that the middle cell can assume. For the case where the internal flow of middle cell is polarized (Fig.S.15A,B) we find that above the $a_c$ transition line (large $a$ values) a persistent CIL configuration (←→→) dominates the phase-diagram (Fig.S.15B,(vi)). As $a$ decreases and $q$ increases, we pass to a region where a full CL configuration (→→→) dominates (Fig.S.15B,(i-v)). In between these phases, and for low values of $a$ and $q$, several multi-stable regions exist (depicted for the low $q$ values in the bifurcation diagrams in Fig.S.15B,(i-v)).

For the case where the local actin flows of the middle cell's edges are activated (Fig.S.15C,D), we find that above the $a_c$ transition line (large $a$ values) the full CIL configuration (← 0 →) dominates the phase-diagram (Fig.S.15C,(v-vi)). As $a$ decreases and $q$ increases, we pass to a region where the anti-CIL configuration (→ 0 ←) dominates (Fig.S.15C,(i-v)). In between these phases, and for low values of $a$ and $q$, several multi-stable regions exist (depicted for the low $q$ values in the bifurcation diagrams in Fig.S.15C,(i-v)). The reason behind the appearance of the anti-CIL configuration for large values of $q$ lies in the propagation of the symmetric fold bifurcation for both of the edge cells along the phase-diagram. As $q$ increase, both branches eventually change a sign (blue and red curves Fig.S.15D).

For the case where the actin polymerization speeds at the middle cell's edges are inhibited (Fig.S.15 E,F), we find that the entire phase-diagram is dominated by a full CIL configuration (← 0 →), except for a multi-stable region of small values of $a$ and $q$ (below the blue-red striped transition line in Fig.S.15E).

### APPENDIX I - EXTENDED STEADY-STATE ANALYSIS OF THE TRIPLET SYSTEM

In this section we present an extend analysis for the steady-state behavior of cell triplets for low and high values of noise. Note here that we count six configurations, i.e the (→ 0 →) configuration introduced at the main text (Fig.7 in main text) is classified as (→←→). We also note that unlike color code in the main text (cell 1/2/3 are blue/teal/red), here the color code is blue/red/magenta for cell 1/2/3.

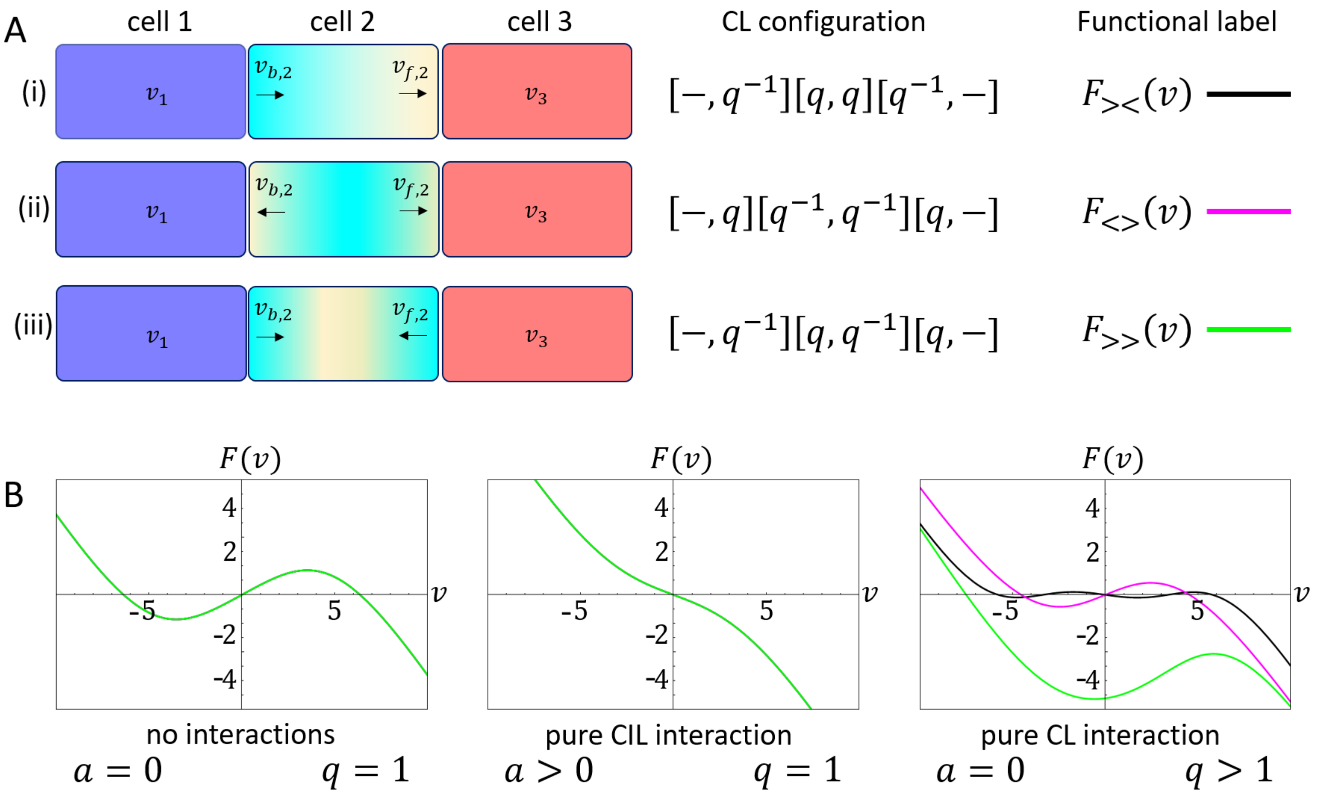


FIG. S.14: The triplet model. A) The possible internal configurations of the middle cell with respect to the CL interaction. i) Polarized state. ii) Non-polarized state in which the two edges are activated. iii) Non-polarized state in which the two edges are inhibited. Blue/Teal/Red indicate cell 1/2/3. B) The functional forms of the steady-state global actin flow in the middle cell (Eqs.S.34-S.36) with respect to cases where there are no interactions (left panel), pure CIL interaction (middle panel), and pure CL interaction (right panel). Purple/Black/Green curves indicate the internal flow with respect to the labels in A.

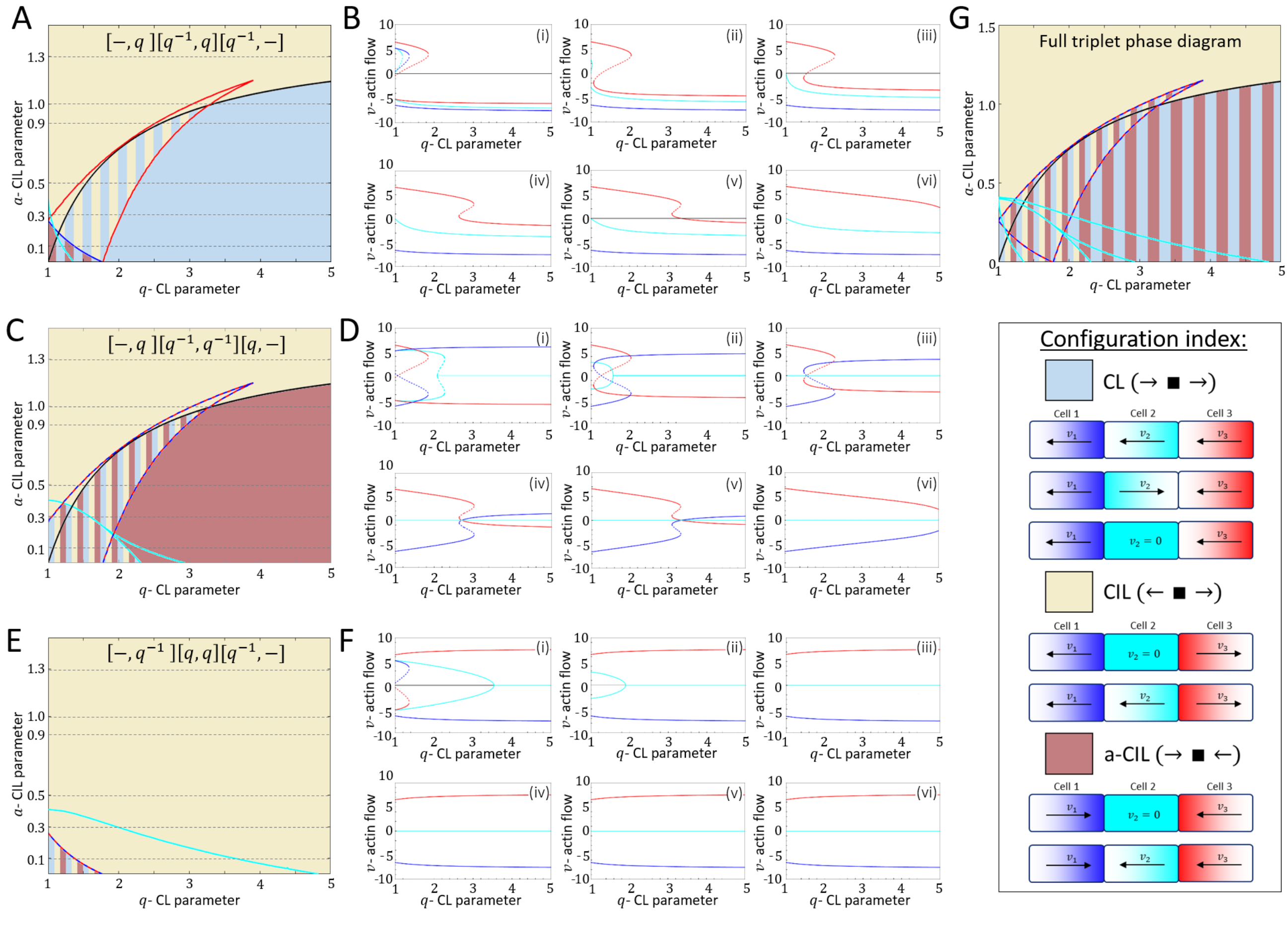


FIG. S.15: A) The $a-q$ phase diagram in which the middle cell is polarized. Blue/Teal/Red curves denote the transition lines for cell 1/2/3. B) Bifurcation diagrams for the cross sections of $a = 0.1, 0.3, 0.5, 0.9, 1.0, 1.3$ in A, denoted by (i-vi) respectively. C) The $a-q$ phase diagram, where the actin polymerization speeds in the edges of the middle cells are activated. D) Bifurcation diagrams for the cross sections of $a = 0.1, 0.3, 0.5, 0.9, 1.0, 1.3$ in C, denoted by (i-vi) respectively. Blue/Red curves denote cell 1/3 or cell 3/1. Teal curve denotes cell 2. E) The $a-q$ phase diagram, where the actin polymerization speeds in the edges of the middle cells are inhibited. F) Bifurcation diagrams for the cross sections of $a = 0.1, 0.3, 0.5, 0.9, 1.0, 1.3$ in (E), denoted by (i-iv) respectively. Blue/Red curves denote cell 1/3 or cell 3/1. Teal curve denotes cell 2. G) The full $a-q$ phase diagram, considering all the configurations of the middle cell. Color index for the phase diagram: Khaki/light blue/brown represent regions of CL/CIL/a-CIL configurations (defined by the edge cells, as depicted in the configuration index). Upper inset denotes the CL interactions between the cells ($q/q^{-1}$ for inhibition/activation). Blue/Teal/Red solid curves in A,C,E denote the transition lines for cell 1/2/3. Striped blue/red curve denote a similar transition line for the edge cells in cases that the behaviors of cells 1 and 3 are symmetric.

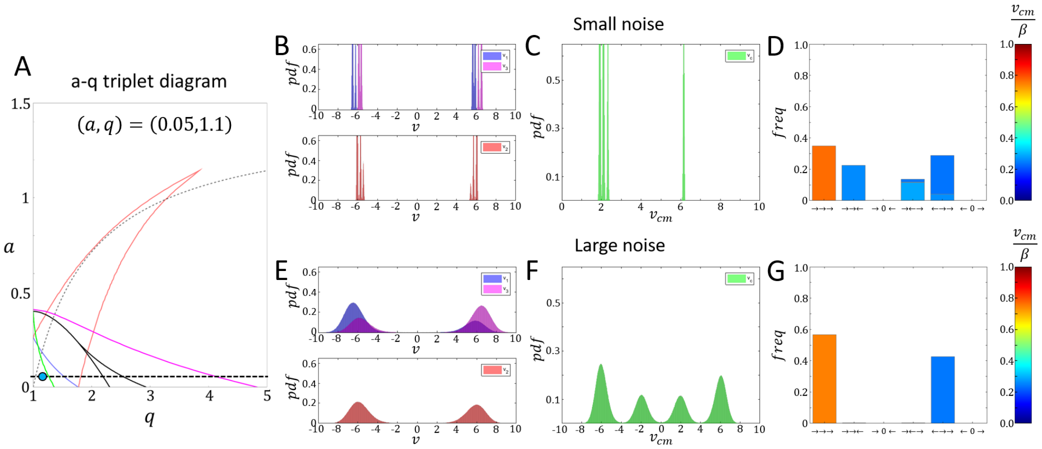


FIG. S.16: Steady-state behavior of a triplet system with $(a, q) = (0.05, 1.1)$. A) The $a$-$q$ triplet phase-diagram indicating the analyzed point (teal circle) along the $a$ cross-section (dashed black horizontal line). B-D) Steady-state dynamics for a small level of noise ($\sigma = 1/32$). E-F) Steady-state dynamics for a large level of noise ($\sigma = 1/\sqrt{2}$). Blue/Red/purple/Green distributions indicate $v_1/v_2/v_3/v_{cm}$ respectively. Bars in D,G indicate the proportions of the configurations with the color code indicating the magnitude of the center-of-mass velocity (normalized by $\beta$).

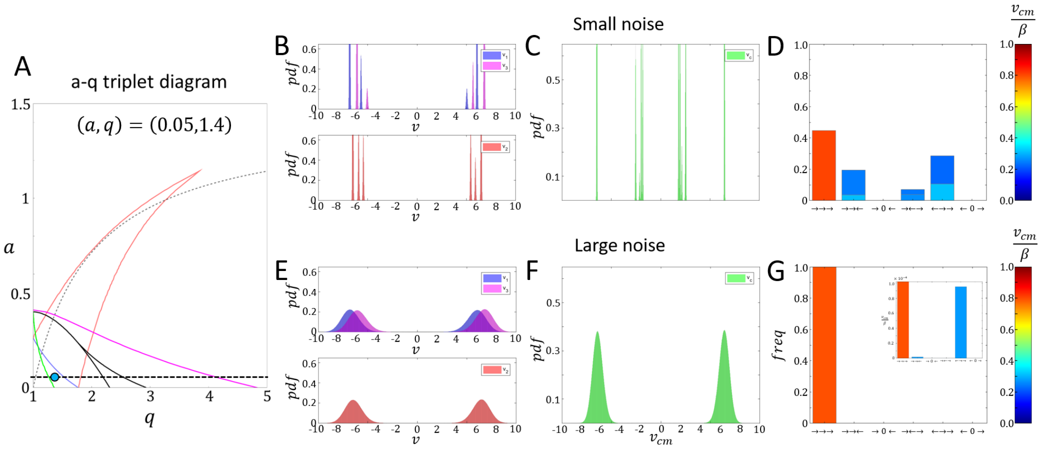


FIG. S.17: Steady-state behavior of a triplet system with $(a, q) = (0.05, 1.4)$. A) The $a$-$q$ triplet phase-diagram indicating the analyzed point (teal circle) along the $a$ cross-section (dashed black horizontal line). B-D) Steady-state dynamics for a small level of noise ($\sigma = 1/32$). E-F) Steady-state dynamics for a large level of noise ($\sigma = 1/\sqrt{2}$). Blue/Red/purple/Green distributions indicate $v_1/v_2/v_3/v_{cm}$ respectively. Bars in D,G indicate the proportions of the configurations with the color code indicating the magnitude of the center-of-mass velocity (normalized by $\beta$).

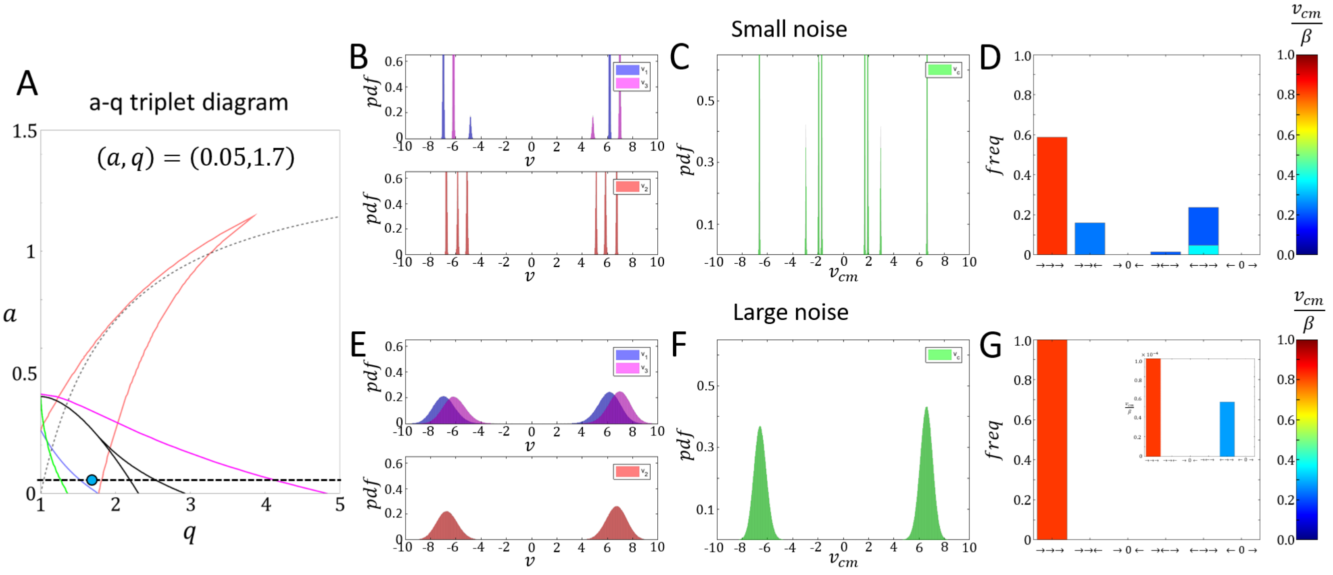


FIG. S.18: Steady-state behavior of a triplet system with $(a, q) = (0.05, 1.7)$. A) The $a$-$q$ triplet phase-diagram indicating the analyzed point (teal circle) along the $a$ cross-section (dashed black horizontal line). B-D) Steady-state dynamics for a small level of noise ($\sigma = 1/32$). E-F) Steady-state dynamics for a large level of noise ($\sigma = 1/\sqrt{2}$). Blue/Red/purple/Green distributions indicate $v_1/v_2/v_3/v_{cm}$ respectively. Bars in D,G indicate the proportions of the configurations with the color code indicating the magnitude of the center-of-mass velocity (normalized by $\beta$).

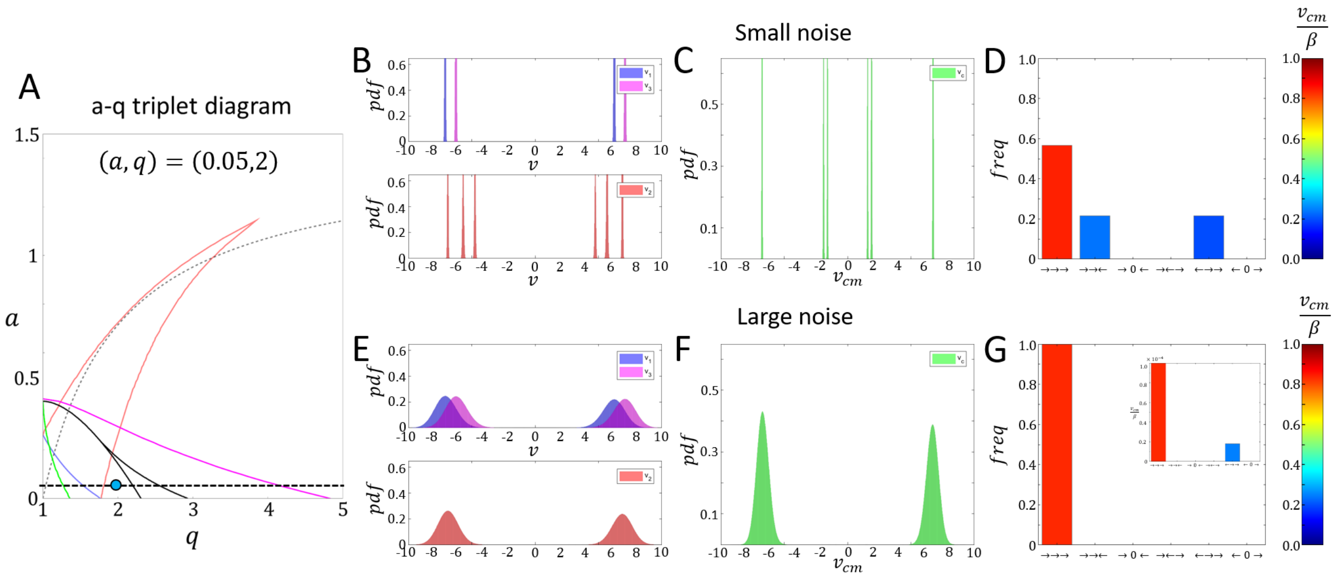


FIG. S.19: Steady-state behavior of a triplet system with $(a, q) = (0.05, 2)$. A) The $a$-$q$ triplet phase-diagram indicating the analyzed point (teal circle) along the $a$ cross-section (dashed black horizontal line). B-D) Steady-state dynamics for a small level of noise ($\sigma = 1/32$). E-F) Steady-state dynamics for a large level of noise ($\sigma = 1/\sqrt{2}$). Blue/Red/purple/Green distributions indicate $v_1/v_2/v_3/v_{cm}$ respectively. Bars in D,G indicate the proportions of the configurations with the color code indicating the magnitude of the center-of-mass velocity (normalized by $\beta$).

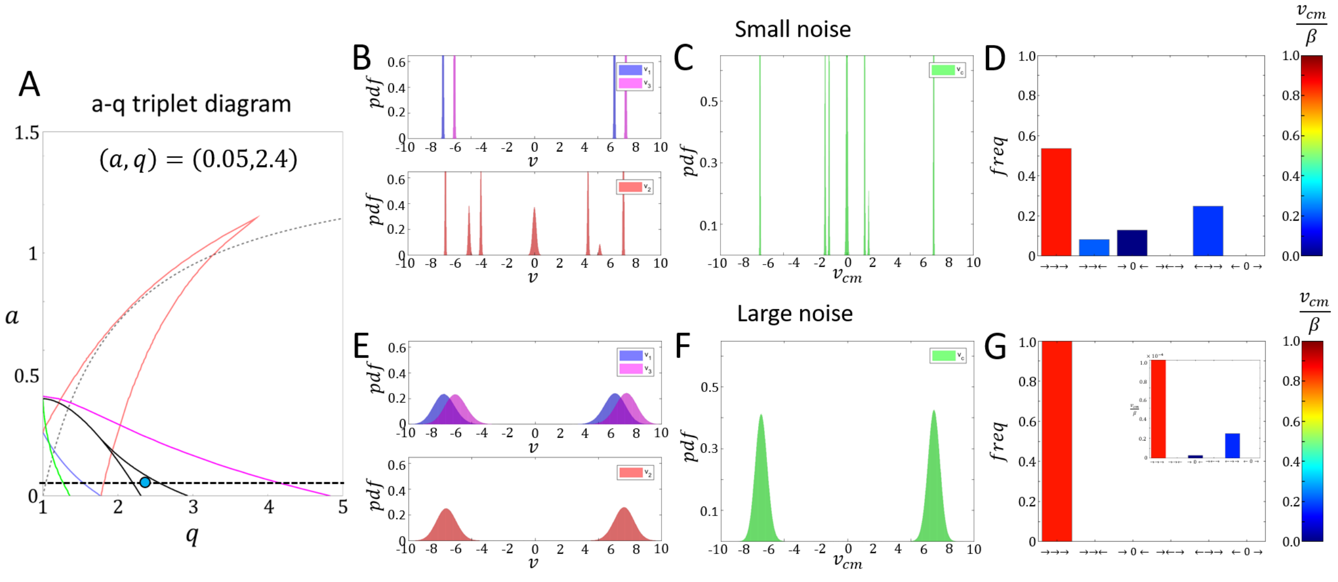


FIG. S.20: Steady-state behavior of a triplet system with $(a, q) = (0.05, 2.4)$. A) The $a$-$q$ triplet phase-diagram indicating the analyzed point (teal circle) along the $a$ cross-section (dashed black horizontal line). B-D) Steady-state dynamics for a small level of noise ($\sigma = 1/32$). E-F) Steady-state dynamics for a large level of noise ($\sigma = 1/\sqrt{2}$). Blue/Red/purple/Green distributions indicate $v_1/v_2/v_3/v_{cm}$ respectively. Bars in D,G indicate the proportions of the configurations with the color code indicating the magnitude of the center-of-mass velocity (normalized by $\beta$).

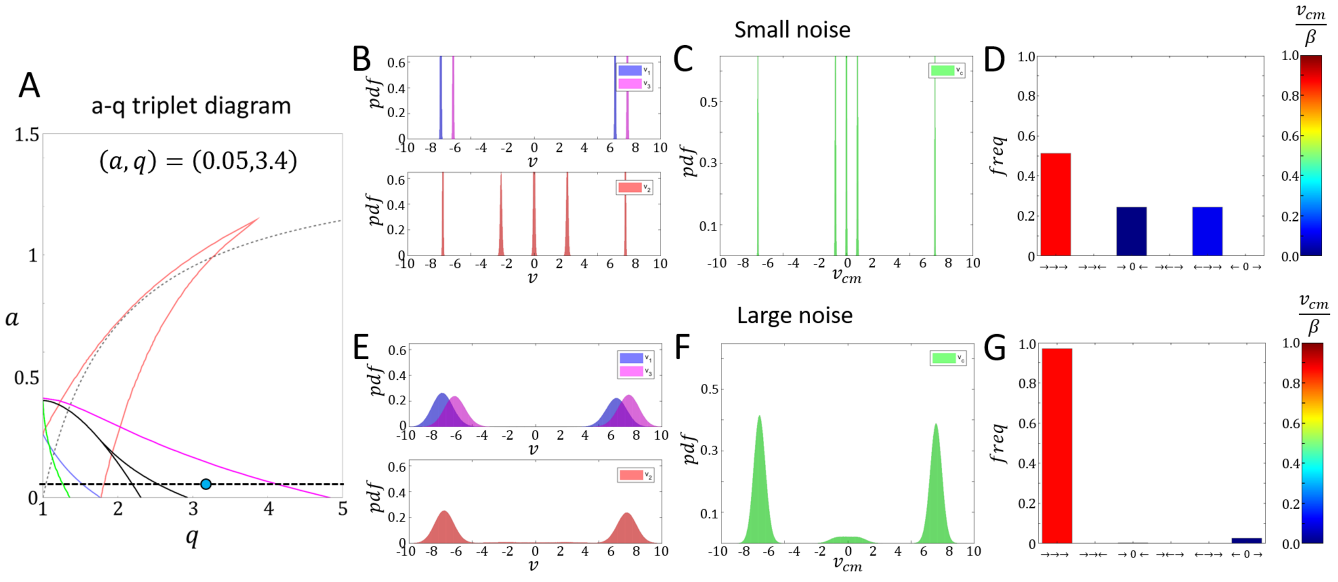


FIG. S.21: Steady-state behavior of a triplet system with $(a, q) = (0.05, 3.4)$. A) The $a$-$q$ triplet phase-diagram indicating the analyzed point (teal circle) along the $a$ cross-section (dashed black horizontal line). B-D) Steady-state dynamics for a small level of noise ($\sigma = 1/32$). E-F) Steady-state dynamics for a large level of noise ($\sigma = 1/\sqrt{2}$). Blue/Red/purple/Green distributions indicate $v_1/v_2/v_3/v_{cm}$ respectively. Bars in D,G indicate the proportions of the configurations with the color code indicating the magnitude of the center-of-mass velocity (normalized by $\beta$).

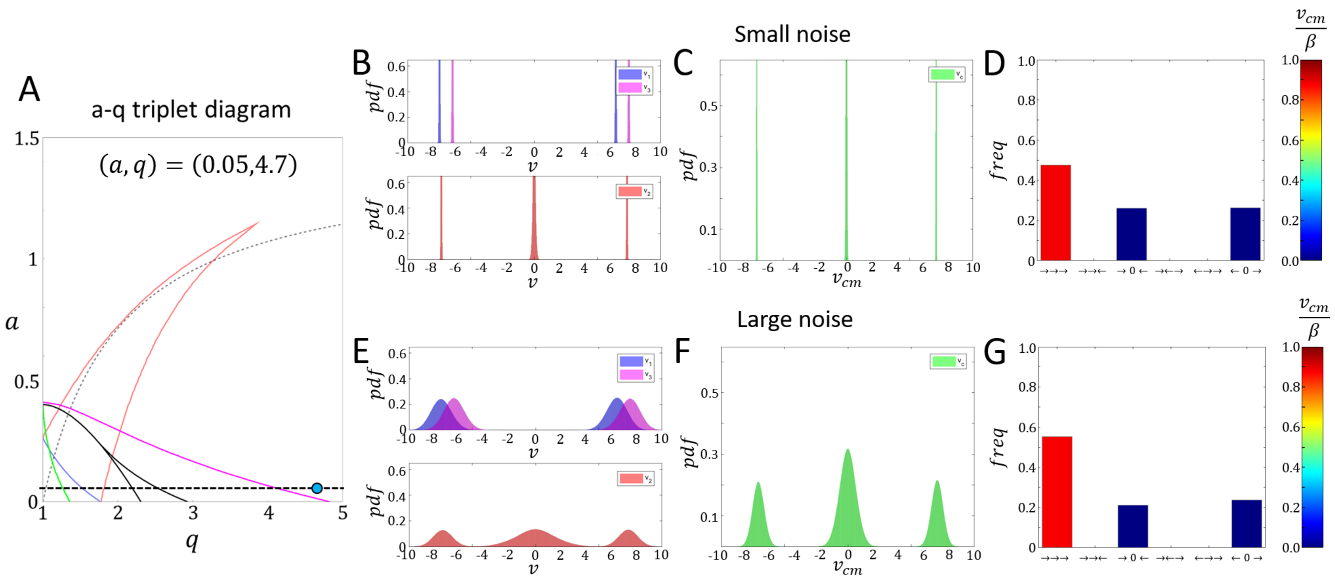


FIG. S.22: Steady-state behavior of a triplet system with $(a, q) = (0.05, 4.7)$. A) The $a$-$q$ triplet phase-diagram indicating the analyzed point (teal circle) along the $a$ cross-section (dashed black horizontal line). B-D) Steady-state dynamics for a small level of noise ($\sigma = 1/32$). E-F) Steady-state dynamics for a large level of noise ($\sigma = 1/\sqrt{2}$). Blue/Red/purple/Green distributions indicate $v_1/v_2/v_3/v_{cm}$ respectively. Bars in D,G indicate the proportions of the configurations with the color code indicating the magnitude of the center-of-mass velocity (normalized by $\beta$).

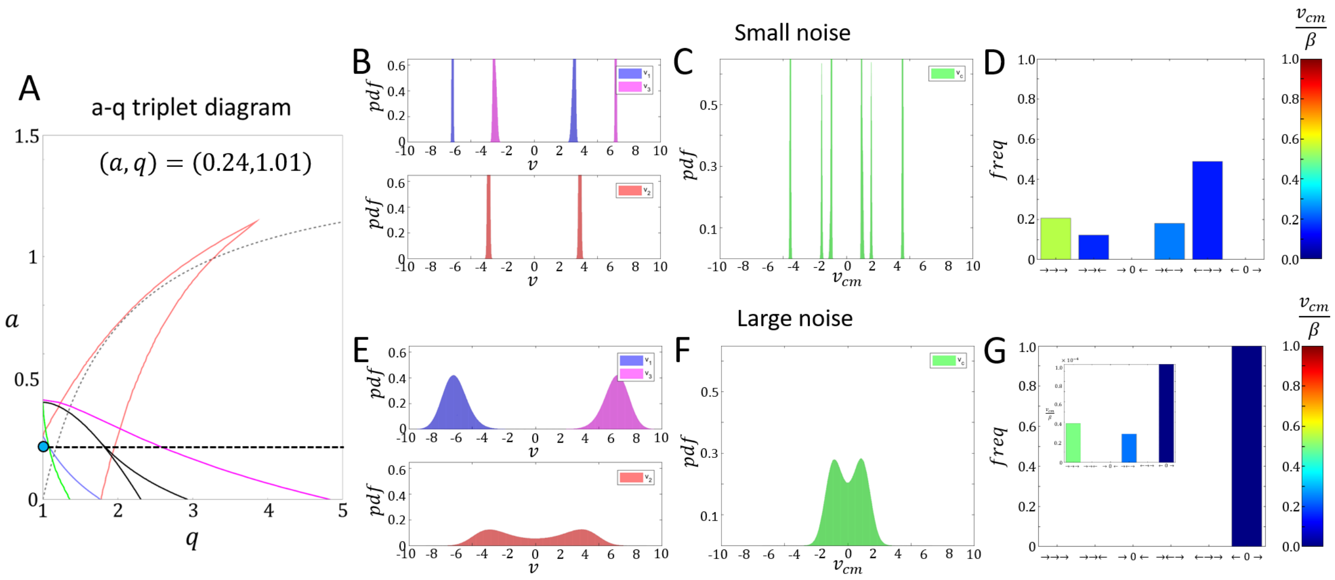


FIG. S.23: Steady-state behavior of a triplet system with $(a, q) = (0.24, 1.01)$. A) The $a$-$q$ triplet phase-diagram indicating the analyzed point (teal circle) along the $a$ cross-section (dashed black horizontal line). B-D) Steady-state dynamics for a small level of noise ($\sigma = 1/32$). E-F) Steady-state dynamics for a large level of noise ($\sigma = 1/\sqrt{2}$). Blue/Red/purple/Green distributions indicate $v_1/v_2/v_3/v_{cm}$ respectively. Bars in D,G indicate the proportions of the configurations with the color code indicating the magnitude of the center-of-mass velocity (normalized by $\beta$).

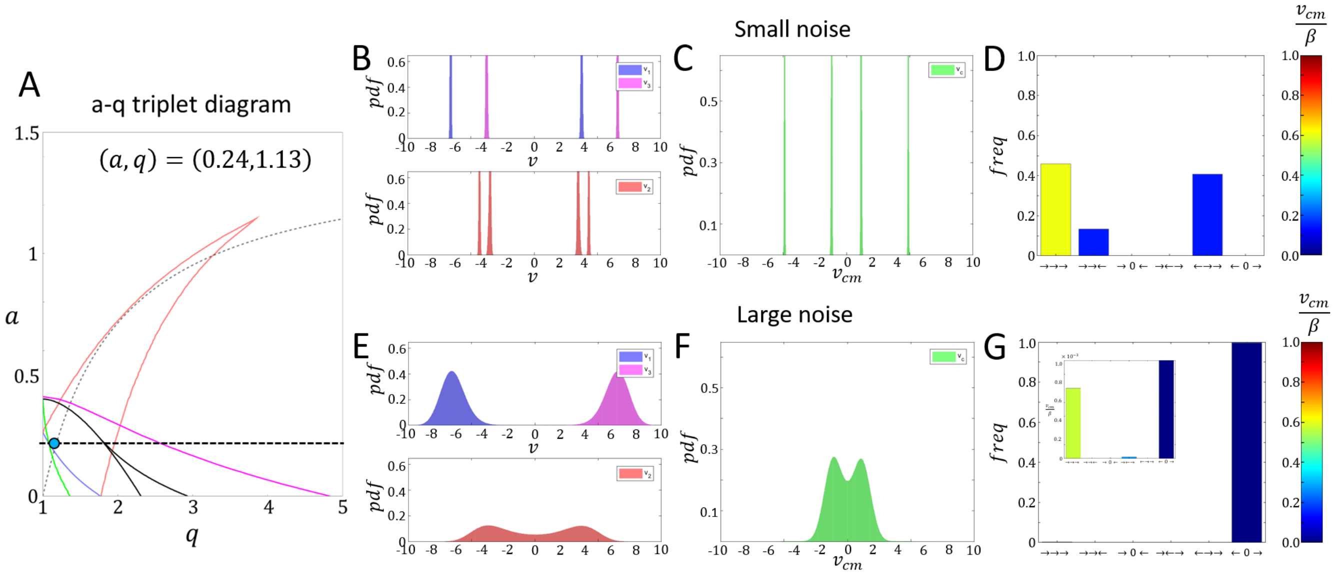


FIG. S.24: Steady-state behavior of a triplet system with $(a, q) = (0.24, 1.13)$. A) The $a$-$q$ triplet phase-diagram indicating the analyzed point (teal circle) along the $a$ cross-section (dashed black horizontal line). B-D) Steady-state dynamics for a small level of noise ($\sigma = 1/32$). E-F) Steady-state dynamics for a large level of noise ($\sigma = 1/\sqrt{2}$). Blue/Red/purple/Green distributions indicate $v_1/v_2/v_3/v_{cm}$ respectively. Bars in D,G indicate the proportions of the configurations with the color code indicating the magnitude of the center-of-mass velocity (normalized by $\beta$).

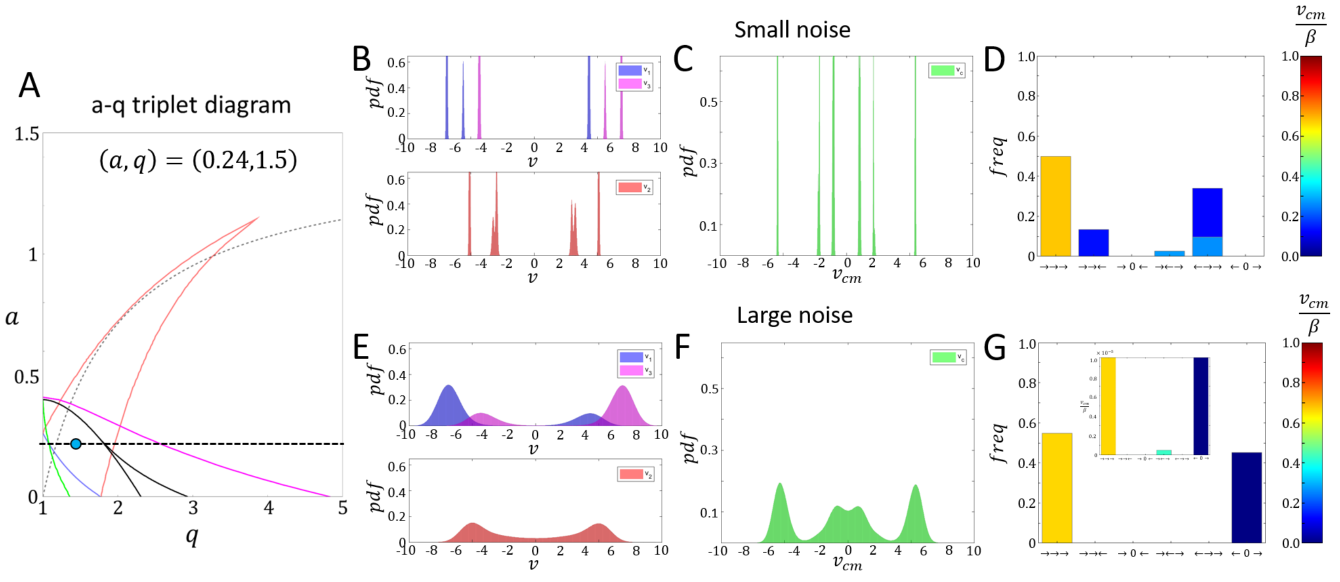


FIG. S.25: Steady-state behavior of a triplet system with $(a, q) = (0.24, 1.5)$. A) The $a$-$q$ triplet phase-diagram indicating the analyzed point (teal circle) along the $a$ cross-section (dashed black horizontal line). B-D) Steady-state dynamics for a small level of noise ($\sigma = 1/32$). E-F) Steady-state dynamics for a large level of noise ($\sigma = 1/\sqrt{2}$). Blue/Red/purple/Green distributions indicate $v_1/v_2/v_3/v_{cm}$ respectively. Bars in D,G indicate the proportions of the configurations with the color code indicating the magnitude of the center-of-mass velocity (normalized by $\beta$).

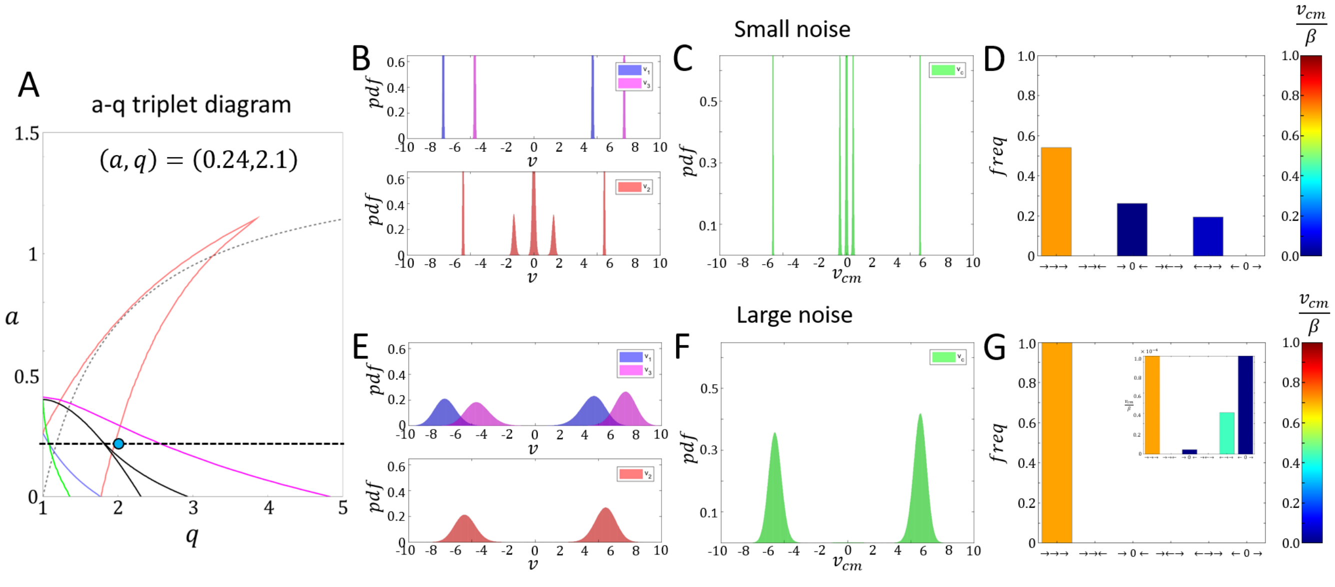


FIG. S.26: Steady-state behavior of a triplet system with $(a, q) = (0.24, 2.1)$. A) The $a$-$q$ triplet phase-diagram indicating the analyzed point (teal circle) along the $a$ cross-section (dashed black horizontal line). B-D) Steady-state dynamics for a small level of noise ($\sigma = 1/32$). E-F) Steady-state dynamics for a large level of noise ($\sigma = 1/\sqrt{2}$). Blue/Red/purple/Green distributions indicate $v_1/v_2/v_3/v_{cm}$ respectively. Bars in D,G indicate the proportions of the configurations with the color code indicating the magnitude of the center-of-mass velocity (normalized by $\beta$).

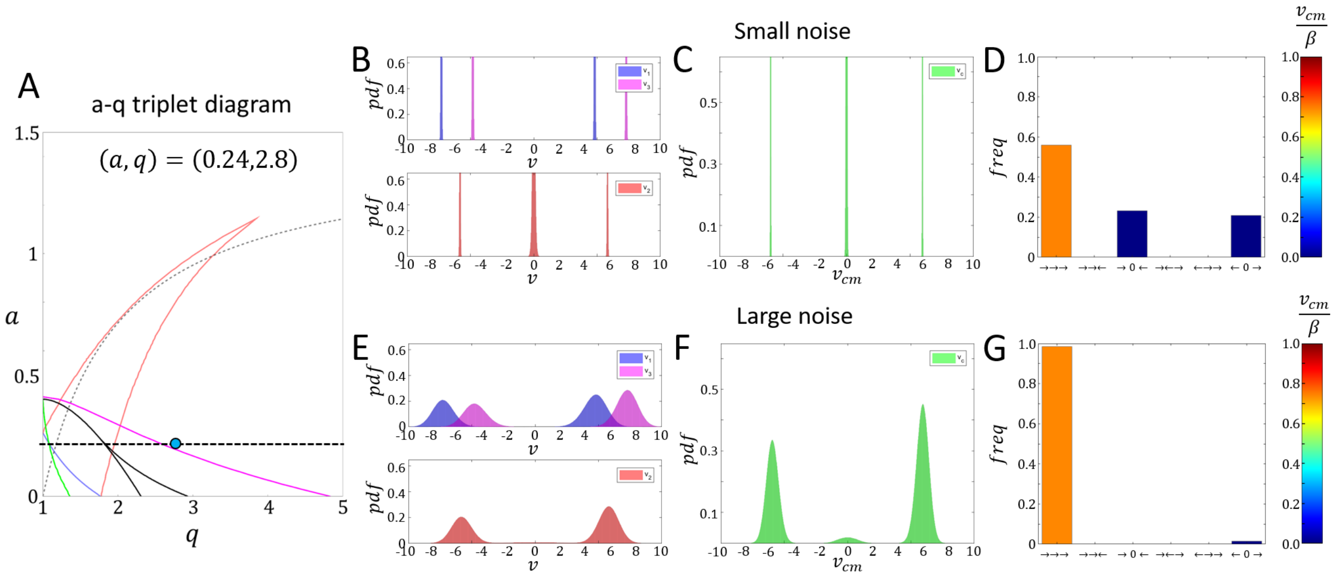


FIG. S.27: Steady-state behavior of a triplet system with $(a, q) = (0.24, 2.8)$. A) The $a$-$q$ triplet phase-diagram indicating the analyzed point (teal circle) along the $a$ cross-section (dashed black horizontal line). B-D) Steady-state dynamics for a small level of noise ($\sigma = 1/32$). E-F) Steady-state dynamics for a large level of noise ($\sigma = 1/\sqrt{2}$). Blue/Red/purple/Green distributions indicate $v_1/v_2/v_3/v_{cm}$ respectively. Bars in D,G indicate the proportions of the configurations with the color code indicating the magnitude of the center-of-mass velocity (normalized by $\beta$).

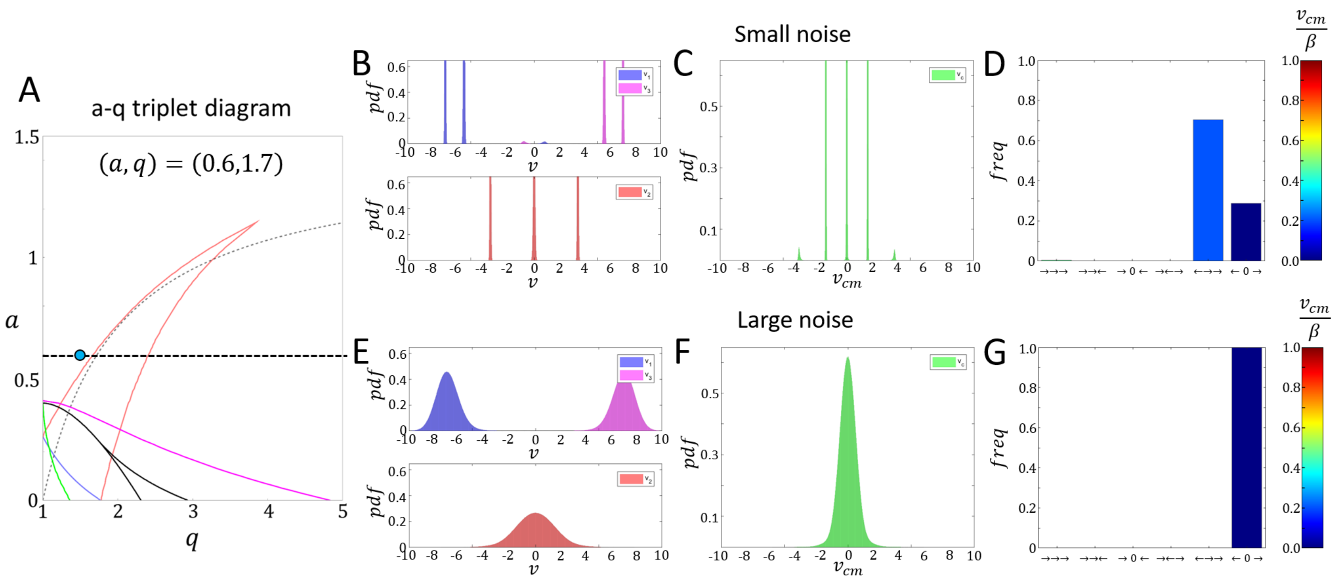


FIG. S.28: Steady-state behavior of a triplet system with $(a, q) = (0.6, 1.7)$. A) The $a$-$q$ triplet phase-diagram indicating the analyzed point (teal circle) along the $a$ cross-section (dashed black horizontal line). B-D) Steady-state dynamics for a small level of noise ($\sigma = 1/32$). E-F) Steady-state dynamics for a large level of noise ($\sigma = 1/\sqrt{2}$). Blue/Red/purple/Green distributions indicate $v_1/v_2/v_3/v_{cm}$ respectively. Bars in D,G indicate the proportions of the configurations with the color code indicating the magnitude of the center-of-mass velocity (normalized by $\beta$).

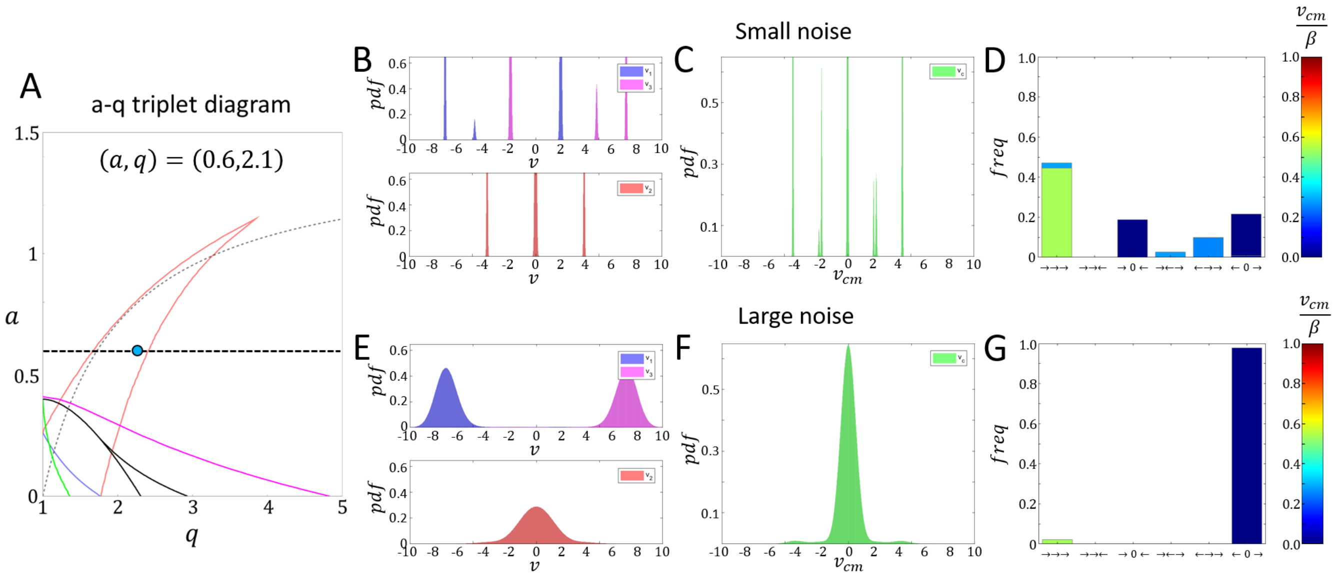


FIG. S.29: Steady-state behavior of a triplet system with $(a, q) = (0.6, 2.1)$. A) The $a$-$q$ triplet phase-diagram indicating the analyzed point (teal circle) along the $a$ cross-section (dashed black horizontal line). B-D) Steady-state dynamics for a small level of noise ($\sigma = 1/32$). E-F) Steady-state dynamics for a large level of noise ($\sigma = 1/\sqrt{2}$). Blue/Red/purple/Green distributions indicate $v_1/v_2/v_3/v_{cm}$ respectively. Bars in D,G indicate the proportions of the configurations with the color code indicating the magnitude of the center-of-mass velocity (normalized by $\beta$).

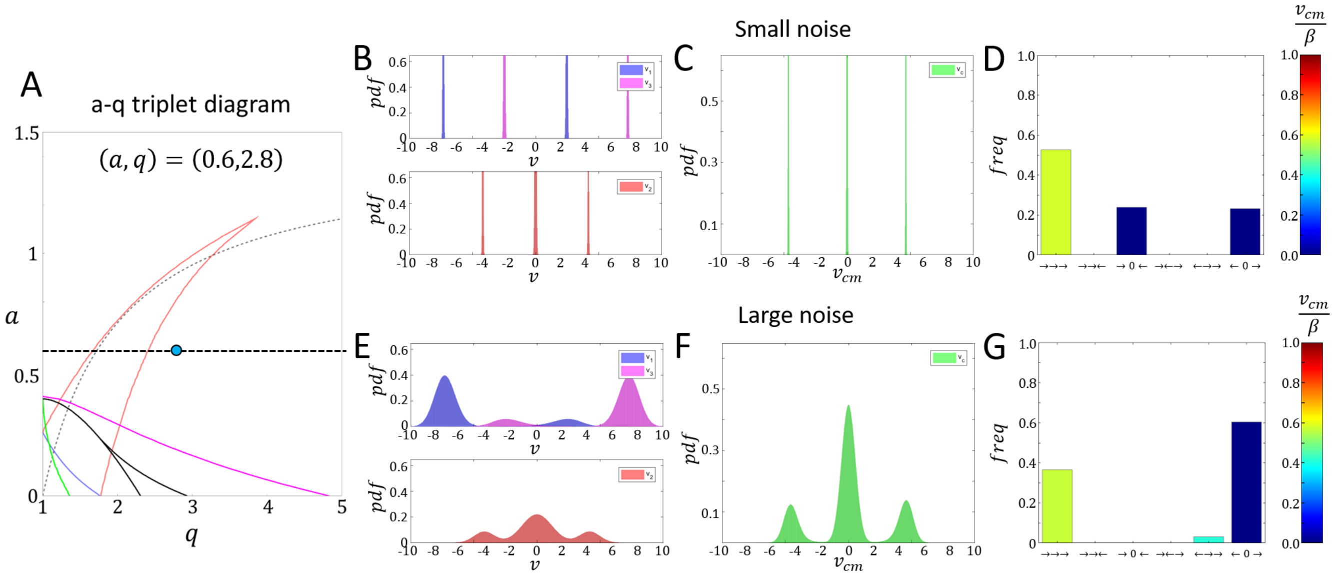


FIG. S.30: Steady-state behavior of a triplet system with $(a, q) = (0.6, 2.8)$. A) The $a$-$q$ triplet phase-diagram indicating the analyzed point (teal circle) along the $a$ cross-section (dashed black horizontal line). B-D) Steady-state dynamics for a small level of noise ($\sigma = 1/32$). E-F) Steady-state dynamics for a large level of noise ($\sigma = 1/\sqrt{2}$). Blue/Red/purple/Green distributions indicate $v_1/v_2/v_3/v_{cm}$ respectively. Bars in D,G indicate the proportions of the configurations with the color code indicating the magnitude of the center-of-mass velocity (normalized by $\beta$).

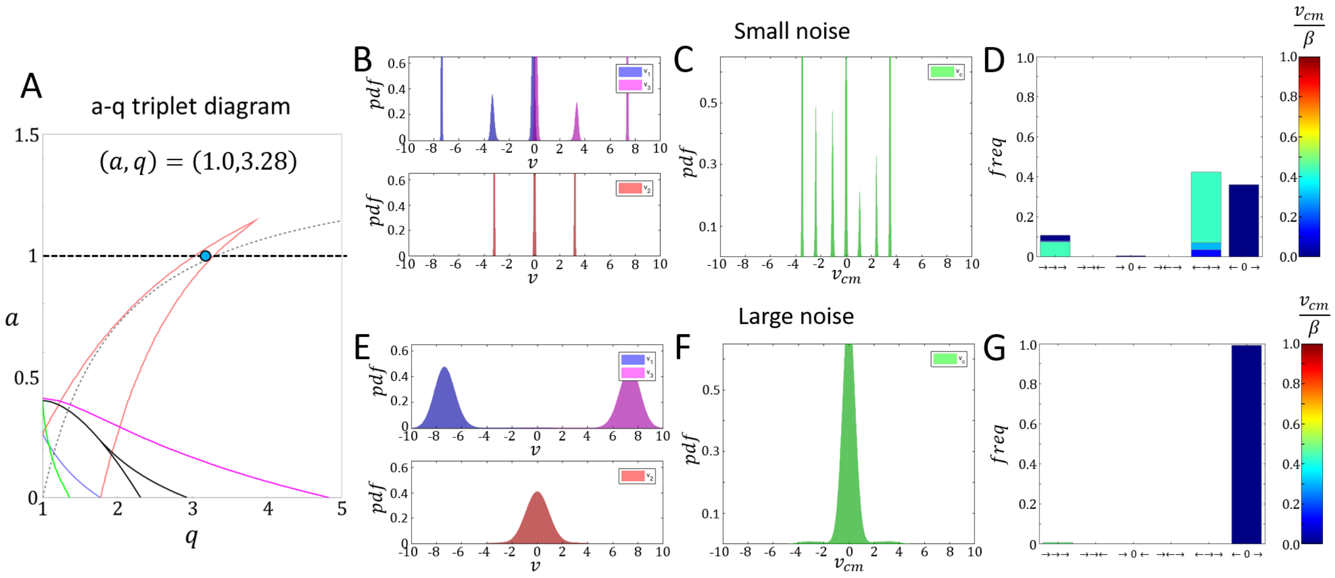


FIG. S.31: Steady-state behavior of a triplet system with $(a, q) = (1, 3.28)$. A) The $a$-$q$ triplet phase-diagram indicating the analyzed point (teal circle) along the $a$ cross-section (dashed black horizontal line). B-D) Steady-state dynamics for a small level of noise ($\sigma = 1/32$). E-F) Steady-state dynamics for a large level of noise ($\sigma = 1/\sqrt{2}$). Blue/Red/purple/Green distributions indicate $v_1/v_2/v_3/v_{cm}$ respectively. Bars in D,G indicate the proportions of the configurations with the color code indicating the magnitude of the center-of-mass velocity (normalized by $\beta$).

### APPENDIX J - CLUSTER DOMAIN DISTRIBUTION FOR A LARGE ALIGNMENT COEFFICIENT ($q$)

In this section we display the domain distributions inside long trains in the parameter regime of large $q$: $(a, q) = (0.05, 5)$. In Fig.S.32 we show the distribution of coherent domain lengths, and typical snapshots of the domains along trains of three different lengths. We find that the large $q$ regime stabilizes the aCIL configurations which act as domain walls, thereby decreasing the size of the coherent domains to $\sim 4$ cells, compared to the average size of $\sim 10$ cells at lower $q$ values (point (3) in Fig.7).

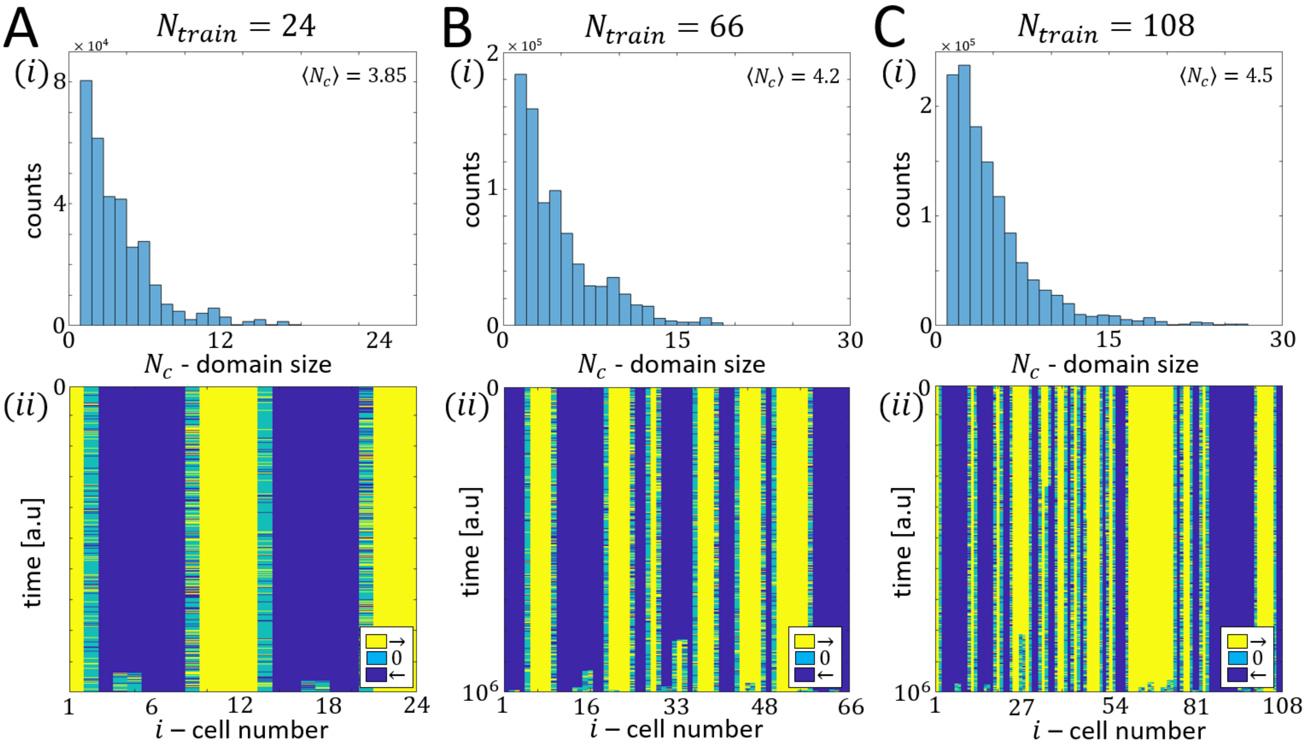


FIG. S.32: J) A-C) Cluster size distribution for trains of size $N = 24/66/108$ in (i) accompanied with an example of the configuration dynamics (ii). Blue/Yellow/Teal colors indicate cells with positive/negative/zero polarization. Parameters: $a = 0.05, q = 5, l = 2.8, d = 4, c = 4$.

### APPENDIX K - RINGS (PERIODIC BOUNDARY CONDITIONS) - EXTENDED ANALYSIS

In this section we present the full heatmaps for cell doublets and triplets on a ring.

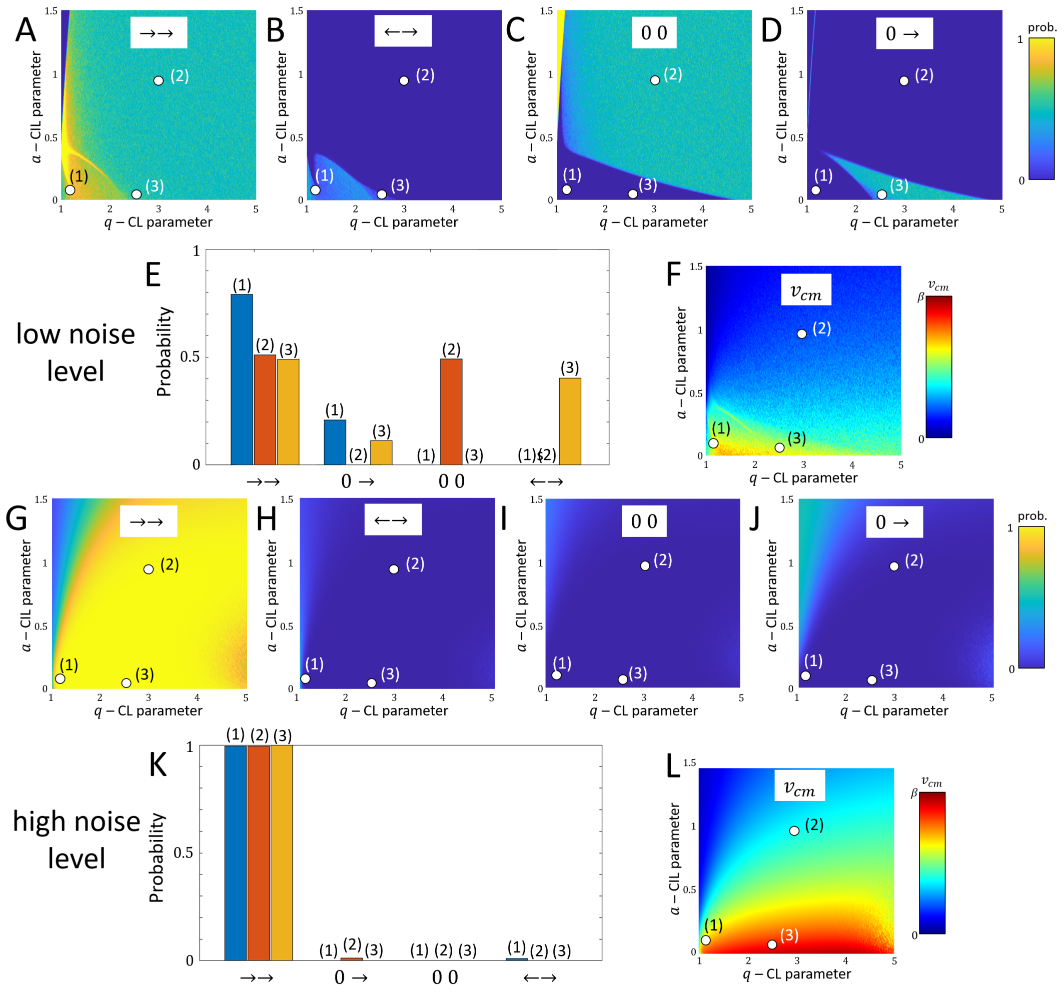


FIG. S.33: Low noise ($\sigma = 1/32$): A-D) Proportion heatmaps for the configurations →→, ←→,00, and 0 →. E) The proportions of the configurations for points (1)-(3) indicated in A-D. F) Center-of-mass velocity heatmap. High noise ($\sigma = 1/\sqrt{2}$): G-L) same as A-F. Parameters: $l = 2.8$,$d = 4$,$c = 4$.

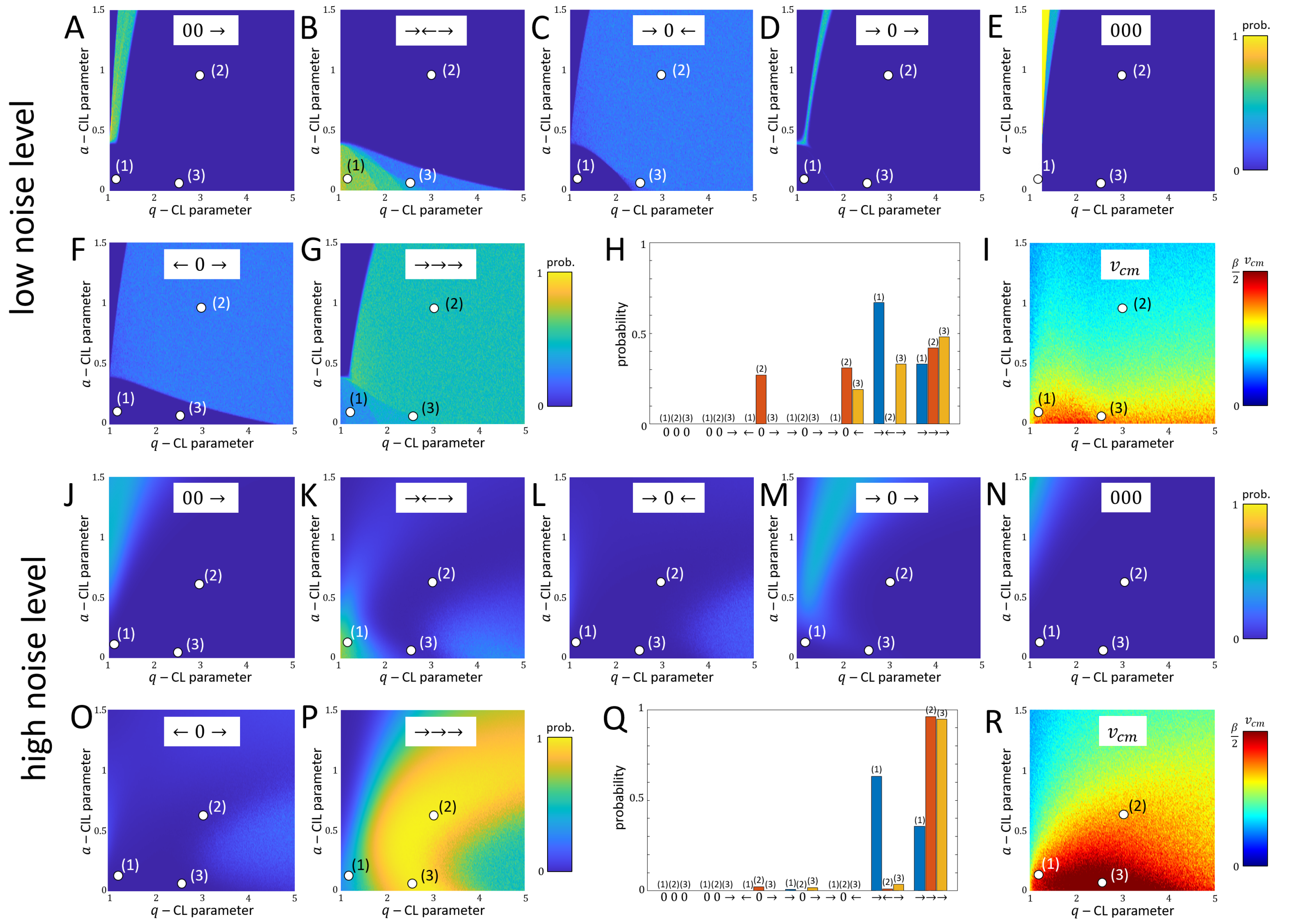


FIG. S.34: Low noise ($\sigma = 1/32$): A-G) Proportion heatmaps for the configurations 0 →→, →←→, → 0 ←,→ 0 →, and 000,← 0 →,→→→. H) Proportions of the configurations for points (1)-(3) in A-G. I) Center-of-mass velocity heatmap. High noise ($\sigma = 1/\sqrt(2)$): J-R) same as A-I. Parameters: $l = 2.8$,$d = 4$,$c = 4$.
